# Gene by environment interaction mouse model reveals a functional role for 5-hydroxymethylcytosine in neurodevelopmental disorders

**DOI:** 10.1101/2021.05.04.441625

**Authors:** Ligia A. Papale, Andy Madrid, Qi Zhang, Kailei Chen, Lara Sak, Sündüz Keleş, Reid S. Alisch

**Affiliations:** Departments of Neurological Surgery, University of Wisconsin, Madison, Wisconsin 53719, USA; Departments of Statistics, Biostatistics, and Medical Informatics, University of Wisconsin, Madison, Wisconsin 53719, USA; Neuroscience training program, University of Wisconsin, Madison, Wisconsin 53719, USA; Department Mathematics and Statistics, University of New Hampshire, Durham, NH 03824, USA

**Keywords:** Developmental brain disorders, Gene by Environment, 5hmC

## Abstract

Mouse knockouts of *Cntnap2* exhibit altered neurodevelopmental behavior and a genome-wide disruption of 5-hydroxymethylcytosine (5hmC). Here we examined whether adult *Cntnap2* heterozygous mice (*Cntnap2*^+/-^, lacking behavioral or neuropathological abnormalities) subjected to a prenatal stress would have disruptions in brain 5hmC levels and exhibit altered behaviors similar to the knockout mice. Adult prenatally stressed *Cntnap2^+/-^* female mice showed repetitive behaviors and altered sociability, similar to the homozygote phenotype. Genomic profiling revealed disruptions in hippocampal and striatal 5hmC levels that were correlated to altered transcript levels of genes linked to these phenotypes (*e.g., Reln*, *Dst*, *Trio* and *Epha5*). Chromatin-immunoprecipitation coupled with high-throughput sequencing and hippocampal nuclear lysate pull-down data indicated that 5hmC abundance alters the binding of the transcription factor CLOCK near the promoters of differentially expressed genes (*e.g., Palld*, *Gigyf1*, and *Fry*), providing a mechanistic role for 5hmC (disruption of transcription factor binding) in gene regulation of developmentally important genes.

## Introduction

Developmental brain disorders usually present years after birth; however, the molecular pathogenesis is thought to arise earlier, either during pregnancy or just after birth. Genetic studies have revealed hundreds of rare risk genes that cumulatively account for less than 25% of neurodevelopmental disorder cases^1,2,3^. The search for environmental factors contributing to the origins of neurodevelopmental disorders has been motivated by their multifactorial genetic heredity and incomplete concordance in monozygotic twins^4^. There is clear evidence that individual differences in the developmental-timing and severity of early-life stressors can predict emotional dispositions later in life and influence the risk for the development of psychopathology^5, 6^. Among the earliest adverse experiences is early-life exposure to maternal depression and anxiety, which confers a lifelong risk for behavioral disturbances in childhood and beyond^7^. This process of “fetal programming” may be mediated, in part, by the impact of the early-life experience on the developing hypothalamic-pituitary-adrenal (HPA) axis^8, 9^. The HPA axis is a dynamic metabolic system that regulates homeostatic mechanisms, such as the ability to respond to stressors in early embryonic development, and is highly sensitive to early life adversities^10^. The HPA axis also governs the activity of sex-specific endocrine mechanisms and responds to stress by altering the neuronal epigenome^11^.

5-hydroxymethylcytosine (5hmC) is an environmentally sensitive epigenetic mark that is highly enriched in neuronal cells^12,13,14,15^ and is associated with the regulation of neuronal activity, suggesting that 5hmC plays an important role in coordinating transcriptional activity in the brain^16^. These findings have prompted investigations into the potential role(s) of 5hmC in mental illness, where it has been linked to developmental brain disorders (*e.g.,* fragile X syndrome, Rett syndrome, and autism) and neurodegenerative diseases (*e.g.,* Huntington’s and Alzheimer’s)^17,18,19,20,21,22,23^. Recently we reported that the *Cntnap2* knockout mouse model of autism exhibits a genome-wide disruption of 5hmC in a significant number of orthologous genes and pathways common to human neurodevelopmental disorders (*N* = 57/232)^24^, supporting an interaction between *Cntnap2* and 5hmC in developmental brain disorders.

The *Cntnap2* (or *Caspr2*) gene encodes a neuronal transmembrane protein member of the neurexin superfamily involved in neuron–glia interactions and clustering of potassium channels in myelinated axons^25, 26^. Humans harboring a homozygous loss of *CNTNAP2* function exhibit severe developmental delays and intellectual disabilities, contributing to a variety of neurodevelopmental disorders such as Gilles de la Tourette syndrome, obsessive-compulsive disorder, cortical dysplasia-focal epilepsy syndrome, autism, schizophrenia, Pitt-Hopkins syndrome, and attention deficit hyperactivity disorder^27^. Mice harboring a homozygous loss of *Cntnap2* function exhibit parallels to the major neuropathological features in autism spectrum disorders, including abnormal social behavior, deficits in communication, and stereotypic repetitive behaviors, cortical dysplasia and focal epilepsy^28^. In addition, these mice exhibit defects in neuronal migration of cortical projection neurons and a reduction in the number of striatal GABAergic interneurons. In contrast, heterozygous *Cntnap2* mutant mice lack any of the behavioral or neuropathological abnormalities found in the homozygous mutant^28^.

Recent evidence in humans does not support a relationship between heterozygous deletions of *CNTNAP2* and neurodevelopmental disorders^29^; however, it remains possible that the combination of heterozygous *CNTNAP2* deletions in a genomic background of increased risk may lead to mental illness^29^. With the onset of *Cntnap2* expression in mice beginning on embryonic day 14, its expression may be sensitive to prenatal (early-life) stressors that alter the neuronal epigenome. Thus, we hypothesized that *Cntnap2* heterozygote mutants exposed to early-life stress would have genome-wide disruptions in 5hmC levels and behavioral deficits, similar to the homozygote mutant. Using a stress paradigm that included seven-days of mild and variable prenatal stress from embryonic day (E) 12 to E18, we provide evidence for a gene by environment model of sex-specific social-related deficits and identify a mechanistic role for 5hmC that contributes to this behavioral outcome.

## Results

### Early-life stressed Cntnap2 heterozygous mice show altered social & repetitive behaviors

To test the long-lasting effects of a prenatal (early-life) environmental stress on the development of altered behaviors, we subjected WT pregnant mice (carrying WT and *Cntnap2^+/-^* pups) to seven-days of variable prenatal stress from embryonic day (E) 12 to E18, which overlaps the onset of *Cntnap2* expression (E14; Methods)^28, 30^. These variable stressors included: 36 hours of constant light, 15 min of fox odor beginning 2 hours after lights on, overnight exposure to novel objects, 5 minutes of restraint stress beginning 2 hours after lights on, overnight novel noise, 12 cage changes during light period, and overnight saturated bedding^31, 32^. These mild stressors were previously published and were selected because they do not include pain or directly influence maternal food intake or weight gain. Importantly, both WT and HET mice were taken from the same litter, providing an ideal internal control for our findings.

Offspring were monitored for consistent maternal/offspring interactions and left undisturbed until weaning day (postnatal day 18). At weaning, same sex offspring of both genotypes were randomly distributed into groups for behavioral or molecular testing and left undisturbed until three months of age. A maximum of three mice from each litter were assigned for behavioral and a maximum of two mice from each litter were assigned for molecular testing (Methods). At three months of age, early-life stressed (ELS) heterozygous *Cntnap2* (ELS-HET) and ELS wild-type (ELS-WT), as well as non-ELS HET (control-HET) and non-ELS WT (control-WT) offspring, were subjected to behavioral testing or sacrificed for molecular analysis. All behavioral groups (ELS and controls for both genotypes and sex, *N* = 9-12 per genotype and sex) were assessed using a variety of behavioral tests for general locomotor activity, anxiety and depression-like levels and social aptitude, including open field, elevated plus maze, light and dark box, forced swim tests and social interaction tests. Only female ELS-HET mice showed altered social and repetitive behaviors in two independent social behavioral tests, the 3-chamber social test and a 10-minute reciprocal social interaction test, respectively. As expected for the highly social strain C57/B6J, female controls (control-WT, ELS-WT, and control-HET) showed a significant preference to be in the chamber and interacting with the cup containing an unfamiliar mouse in the 3-chamber social test. In contrast, female ELS-HET mice mirror the homozygous *Cntnap2* mutants by not showing a significant preference (differential in mean time spent in each chamber was 65.3 seconds (s) for control-WT, 80.3s for ELS-WT, 102.4s for control-HET, and 27.8s for ELS-HET; Two-way Anova, *P*-value < 0.05 for controls only; **Figure 1A-B; Supplemental Fig. 1A&D**)^28^. Similarly, the time spent interacting with the unfamiliar mouse instead of the empty cup during the 3-chamber test was significant for the controls but not for the ELS-HET female mice (differential in mean time spent interacting in each chamber was 50.2s for control-WT, 53.6s for ELS-WT, 65.4s for control-HET, 16.1s for ELS-HET; Two-way Anova, *P*-value < 0.05 for controls only; **Figure 1C-D**). The mice were then subjected to an additional social interaction test that confirmed the female ELS-HET mice spent significantly more time grooming/digging than interacting with an unfamiliar mouse compared to all controls (total time spent grooming/digging among pairs of mice matched for treatment, genotype, and sex are 96.4s for control-WT, 74.4s for ELS-WT, 101.2s for control-HET, and 160.8s for ELS-HET; One-way Anova, *P*-value < 0.05, **Figure 1E**). This repetitive grooming/digging behavior phenotype was only present in a social setting, as it was not observed in mice allowed to explore a cage alone for 10 min (total time spent grooming/digging during a 10 minute cage exploratory test are 64.5s for control-WT, 52.2s for ELS-WT, 59.3s for control-HET, and 44.2s for ELS-HET; Two-way Anova, *P*-value > 0.05, **Table 1**). Deficits in these 3 social-related measures were reproducible in an independent cohort of mice (*N* = 8-10 per treatment, sex, and genotype, **Supplemental Fig. 1)**. Notably, male mice (control-WT, control-HET, ELS-WT, and ELS-HET) did not show any altered social or repetitive behavior phenotypes (**Supplemental Fig. 2**). In addition, all behavioral groups showed no significant differences in general locomotor activity and anxiety and depression-like levels (*P*-value > 0.05; **Table 1 and Supplemental Table 1**). Together, these data support altered social and repetitive behaviors in female ELS-HET mice, revealing a sex-specific gene by environment (GxE) interaction effect on behavior. Since homozygous mutants show similar altered social and repetitive-like behaviors^28^ and a genome-wide disruptions in 5hmC^24^, we next sought to examine the 5hmC profile in this GxE interaction model.

**Figure 1:**
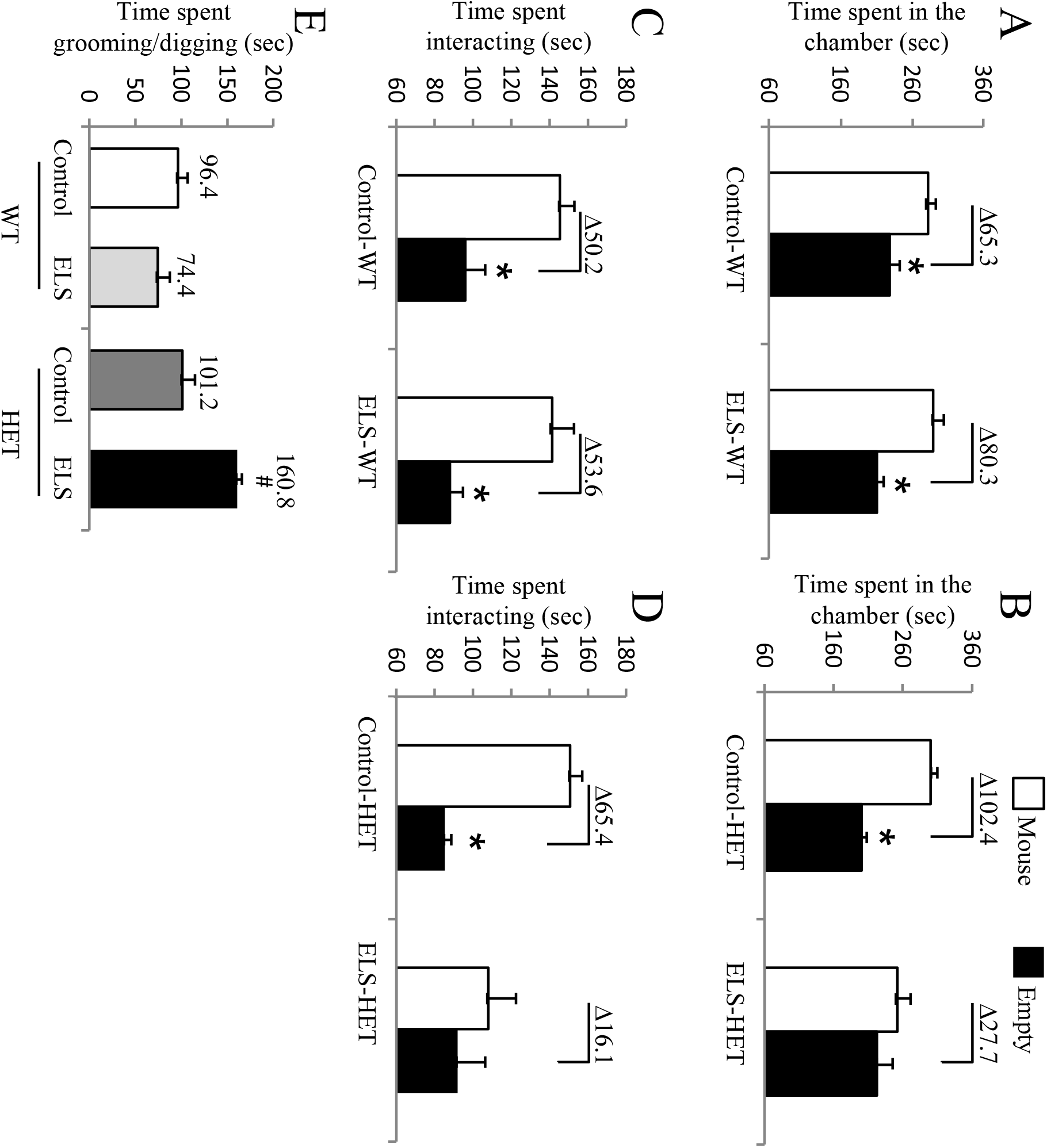
Results from social behavioral tests in female offspring. (A-D) 3-chamber social interaction test: (A-B) time spent in the chamber with either an unfamiliar mouse inside a cup (mouse) or with an empty cup (empty). Data analysis details: (A) two way ANOVA for <u>control-WT versus ELS-WT groups</u>: effect of time spent in either chamber: *F*_(1,42)_ = 28.5, *P*-value < 0.001. (B) two way ANOVA <u>for control-HET versus ELS-HET groups:</u> effect of time spent in either chamber: *F*_(1,32)_ = 16.2, *P*-value < 0.001 and interaction between time spent in either chamber and treatment: *F*_(1,32)_ = 5.3, *P*-value = 0.002. (C-D) Time spent interacting with either an unfamiliar mouse inside a cup (mouse) or with an empty cup (empty). Data analysis details: (C) two way ANOVA for <u>control-WT versus ELS-WT groups:</u> effect of time spent interacting: *F*_(1,42)_ = 29, *P*-value < 0.001. (D) two way ANOVA for <u>control-HET versus ELS-HET groups</u>: effect of time spent interacting in either chamber: *F*_(1,32)_ = 9.9, *P*-value = 0.003). All asterisks denote *P*-value ≤ 0.05, two-way ANOVA with Bonferroni multiple comparison test determined post hoc significance. (E) Ten-minute reciprocal social interaction test in female offspring. One-way ANOVA detected effect of groups on time spent grooming/digging during a 10-minute reciprocal social interaction test (*F*_(3,16)_ = 10.9, *P-*value < 0.001). Hashtag (#) denotes significance for time spent grooming/digging between ELS-HET in comparison to Control-WT, Control-HET, and ELS-WT (*P*-value < 0.05). *N* = 9-10 per treatment and genotype.

**Table 1:**
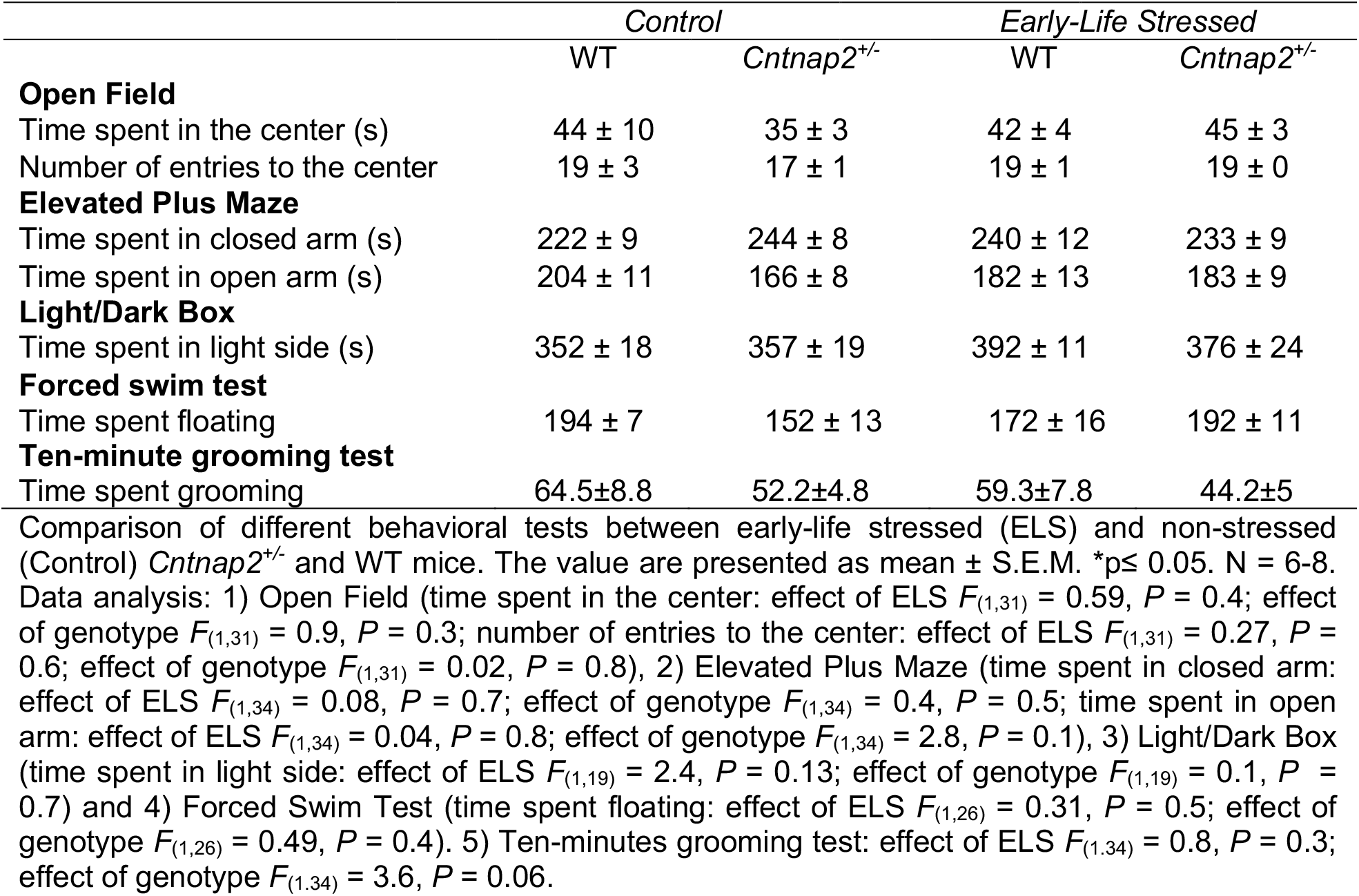
Summary of behavioral tests in female offspring

### Long-lasting disruption of 5hmC in the female GxE interaction model

To determine the effect of this GxE interaction on the genome-wide distribution of 5hmC, we employed an established chemical labeling and affinity purification method, coupled with high-throughput sequencing technology^12, 24, 33, 34^. Age-matched ELS-HET and control HET, as well as ELS-WT and control-WT female offspring (*N* = 6/group) were sacrificed at 3 months of age as independent biological replicates. 5hmC containing DNA sequences were enriched from pools (*N* = 2) of hippocampal tissues, a brain region involved in the pathology of social behavior with high expression of the *Cntnap2* gene since embryonic day 14 and previously shown to be correlated with altered sociability in the *Cntnap2*^-/-^ mice^35^. High-throughput sequencing resulted in a range of ∼25-43 million uniquely mapped reads from each pool of biological replicates (**Supplemental Table 2**, methods). These data showed no visible differences among the chromosomes, except depletion on chromosomes X, which is consistent with previous observations^12, 24, 34^, and an overall equal distribution of 5hmC levels in both genotypes and in both conditions on defined genomic structures and repetitive elements. (**Supplemental Fig. 3**).

### Identification and characterization of differentially hydroxymethylated regions (DhMRs) in the female GxE interaction model

To identify distinct 5hmC distribution patterns following early-life stress throughout the genome of both *Cntnap2* heterozygous and WT female mice, we characterized differentially hydroxymethylated regions (DhMRs) between: 1) ELS-HET and control-HET groups, to investigate the long lasting effects of the GxE interaction on 5hmC levels (HET-DhMRs); as well as 2) ELS-WT and control-WT groups, to determine the long-lasting effects of early-life stress on 5hmC levels (WT-DhMRs). A total of 1,381 ELS-HET increases in hydroxymethylation (ELS-HET-specific 5hmC) and 1,231 ELS-HET decreases in hydroxymethylation (control-HET-specific 5hmC) were found and these loci were distributed across all chromosomes (**Figure 2A**; Dataset 1; methods). In contrast, only a total of 735 significant ELS-WT increases in hydroxymethylation (ELS-WT-specific 5hmC) and 710 significant ELS-WT decreases in hydroxymethylation (control-WT-specific 5hmC) were found and these loci were distributed across all chromosomes (**Supplemental Fig. 4A**; Dataset 1; methods). These data indicate that the GxE interaction results in nearly twice the number of changes in 5hmC levels, but also suggest that early-life stress alone is sufficient to cause stable changes in 5hmC levels. Since specific regions of the genome are differentially methylated based on the biological functions of the genes contained within each region, we next determined if GxE and ELS-induced DhMRs are enriched or depleted on certain chromosomes using a binomial test of all detected 5hmC peaks as the background (Methods). This analysis revealed that DhMRs were not significantly enriched or depleted on any chromosomes (binomial *P*-value < 0.05; **Figure 2A; Supplemental Fig. 4A**).

**Figure 2:**
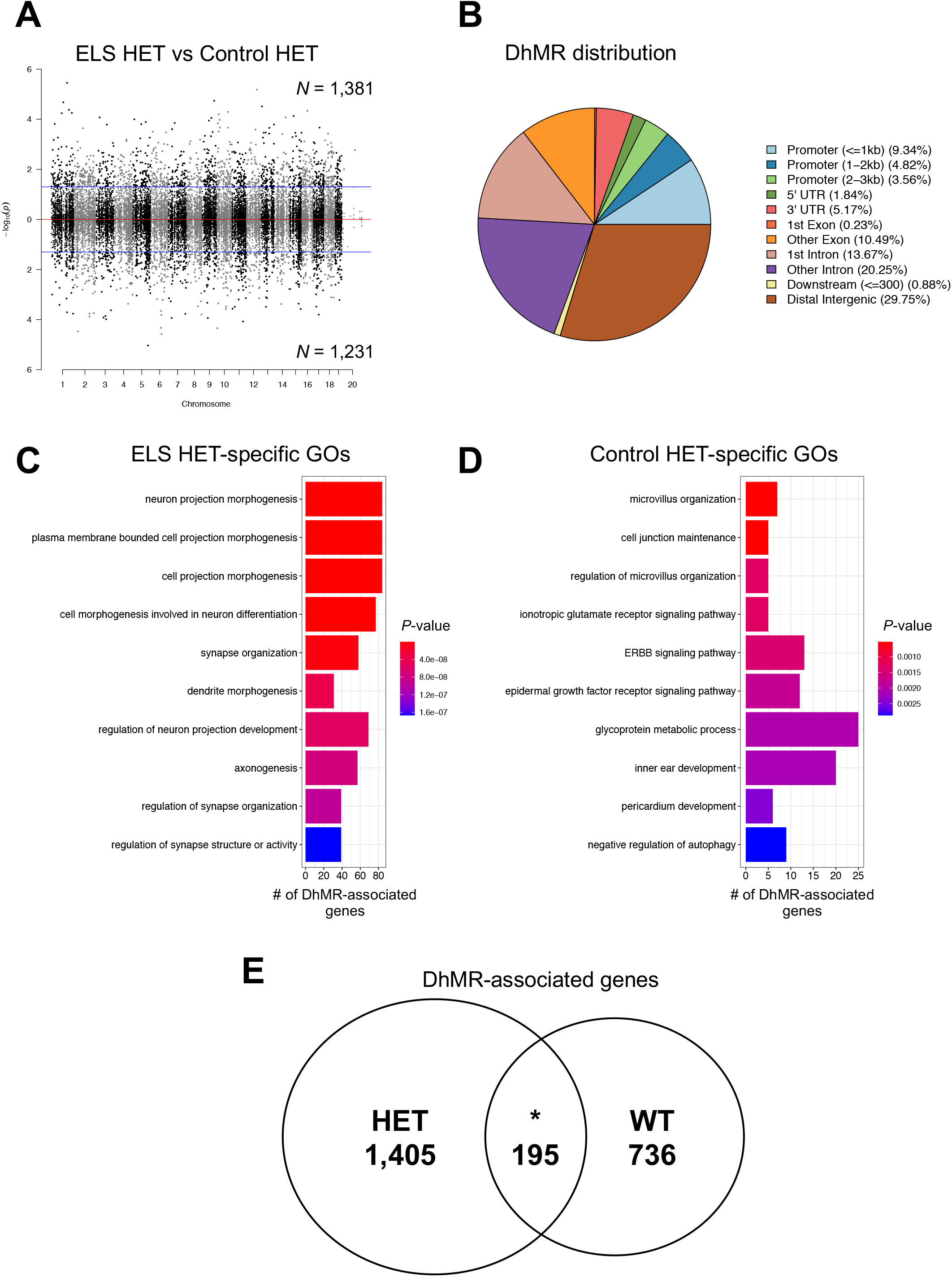
Distribution of hippocampal HET DhMRs. (A) A Manhattan plot shows the distribution of 5hmC sequenced data across the genome (x-axis; alternating black/grey for alternating chromosomes) and the level of significance (-log10(*P*-value)) along the y-axis. Dots above the top blue line represent ELS-HET-specific (hyper)-DhMRs, while dots below the bottom blue line represent Control-HET-specific (hypo)-DhMRs (*P*-value < 0.05). (B) A pie chart displays the proportion of DhMRs across standard genomic structures. (C-D) Bar plot showing the top ten significantly over-represented ontological terms (y-axis) based on gene ontological analysis of ELS-HET-specific DhMR-associated genes (C) and Control-HET-specific DhMR-associated genes (D). The number of DhMR-associated genes linked to the ontological terms are displayed on the x-axis. Bar color is based on *P*-value. (E) Venn diagram showing the overlap of HET-DhMR-associated genes and WT-DhMR-associated genes.

We proceeded to define the genomic features associated with the GxE interaction by annotating HET-DhMRs to overlapping gene structures or to intergenic regions if more than 10 kilobases (kb) away from any gene structure. The overall distribution of these data indicated that the largest fractions of HET-DhMRs were intronic (∼34%) or in distal intergenic regions (∼30%) (**Figure 2B**). Binomial testing revealed that the HET-DhMRs were significantly depleted in exons and promoter regions (**Supplemental Fig. 5A**; binomial *P*-value < 0.05). While the distribution of WT-DhMRs also revealed structures with significant fluctuations, the genomic location of the WT-DhMR fluctuations were unique to those found for the HET-DhMRs (**Supplemental Figs. 4B and 5B** Together, these data indicate that the GxE interaction- and stress-alone-induced DhMRs are unique and not randomly distributed throughout the genome.

Finally, DNA hypermethylation on repetitive elements is believed to play a critical role in maintaining genomic stability^36^. To investigate the genome-wide disruptions of 5hmC on repetitive elements, we aligned the total 5hmC reads and DhMRs to RepeatMasker and the segmental duplication tracks of NCBI37v1/mm9 and found that the largest fraction of HET-DhMRs (∼70%) was in the SINE and simple repeat regions of the genome (**Supplemental Fig. 5C**). The WT-DhMRs were enriched in LTR repeats and were specifically enriched in DNA repeats and SINEs (ELS-WT- and Control-WT-specific, respectively; **Supplemental Fig. 5D**; binomial *P*-value < 0.05). These data suggest a long-term stability of 5hmC at the majority of repeat sequences in female mice following early-life stress, which could either imply stabilization of 5mC in repetitive elements and appear as hypermethylation, a response previously observed in response to stress^37^, or hypomethylation at these sites (removing the ability to generate 5hmC), a response previously observed in PTSD patients (but not controls) following repeated stress^38^.

### Annotation of GxE interaction DhMRs to genes reveals known and potentially novel stress and neurodevelopmental-related genes

To determine the genes and pathways affected by the long-lasting molecular effects of a GxE interaction on female mice, we annotated the HET-DhMRs to genes and found 895 and 770 genes that contained ELS-HET- and control-HET-specific 5hmC levels, respectively. One-hundred and ninety-five genes were found to have more than one DhMR, leading to a total of 1,600 unique genes with disruptions in 5hmC levels (Dataset 1). While sixty-five genes were identified to have both ELS-HET- and control-HET-specific 5hmC, DhMRs residing on the same gene were at distinct genomic loci (average >50kb away from each other), suggesting they have unique molecular roles (*e.g.,* gene regulation) following a GxE interaction. A significant number of HET-DhMR-associated genes were found in well-known stress genes such as *Homer 1*^39^ *and Ncam1*^40^ (524/1600 χ^2^ test *P*-value < 0.0001). Moreover, several of the HET-DhMR-associated genes were previously linked with developmental brain disorders including genes that encode synaptic proteins/receptors (*e.g.,* glutamate, GABAergic, calcium channels, potassium channels, sodium channels, as well as neurexin and neuroligin transmembrane proteins). To examine whether the DhMR-associated genes were enriched for neurodevelopmental-related genes, we compared them to a list of developmental brain disorder genes (*N* = 232 genes; methods)^41^. This comparison revealed that a significant number of orthologs (*N* = 56 of 232; hypergeometric test *P*-value < 0.01) harbor HET-DhMRs, suggesting that 5hmC has a molecular role in the disruption of neurodevelopment following a GxE interaction. Indeed, this finding is similar to the DhMR-associated genes found in the *Cntnap2* homozygous mouse^24^. The HET-DhMR-associated genes also included several genes known to function in epigenetic pathways, including *Dnmt3a*, *Tet2*, and *Hdac4.* To further examine the biological significance of the HET DhMRs-associated genes, we performed separate gene ontological (GO) analyses of the ELS- and control-specific HET DhMR-associated genes (*N* = 1,201 and 1,129, respectively) and found a significant enrichment of neuronal ontological terms only among the ELS-specific HET DhMR-associated genes, which included regulation of synaptic organization, axon development, and neuron projection morphogenesis (*P*-value < 0.05 & FC > 1.5 for GO enrichment tests; **Figure 2C**; Dataset 2). In contrast, the control-specific HET DhMR-associated genes were related to organ-related developmental and morphological processes (**Figure 2D**; Dataset 2). Together, increased 5hmC in the GxE model results in disruptions of genes that encode synaptic proteins/receptors and signaling pathways, supporting a link to brain-related disorders.

Genomic annotation of the WT*-*DhMRs to genes revealed that 510 and 450 genes contained ELS-WT- and control-WT-specific 5hmC, respectively. Sixty genes were found to have more than one DhMR, leading to a total of 931 unique genes with disruptions in 5hmC levels (Dataset 1). While twenty-eight genes were identified to have both ELS-WT- and control-WT-specific 5hmC, WT-DhMRs residing on the same gene were at distinct genomic loci (average >50kb away from each other), suggesting that when found proximal to the same gene WT-DhMRs likely have unique molecular roles (*e.g.,* gene regulation) following early-life stress. Notably, nearly 80% of WT-DhMR-associated genes were unique from the HET-DhMR-associated genes (**Figure 2E**), indicating that the majority of hydroxymethylation changes were specific to genotype.

A significant number of WT DhMR-associated genes were previously linked to stress, including *Dnmt1*, *Pten*, and *Fkbp5*^42,43,44^ (312/931; hypergeometric test *P*-value < 0.01). Moreover, a significant number of orthologs, best known for their role in human neurodevelopmental disorders, have disruptions in 5hmC levels (35/233; hypergeometric *P-*value < 0.01; *e.g., Aut2*, *Nckap5*, *Setbp1*^24, 41^), further implicating stress-alone in the pathogenesis of neurodevelopmental disorders. Finally, WT DhMR-associated genes included *Dnmt1* from the epigenetic pathway. Annotation of the ELS- and control-specific WT DhMR-associated genes (*N* = 697 and 666 genes, respectively) to GO terms found a significant overrepresentation of terms connected with metabolism/catabolism and limb development/morphogenesis (**Supplemental Fig. 4C and D**; Dataset 2). Together, these data indicate that early-life stress alone disrupts 5hmC on biologically relevant genes and pathways.

### Confirmation of long-lasting disruptions of 5hmC in the GxE interaction model

As an initial confirmation of genome-wide disruption of 5hmC following early-life stress, we profiled 5hmC levels in an independent cohort of mice and another brain region associated with social and stress-related behaviors (striatum) that was recently to shown may contribute to pathways of risk in female autism spectrum disorders^45^. Age-matched ELS-HET and control-HET, as well as ELS-WT and control-WT female mice (*N* = 3/group) were sacrificed at 3 months of age as independent biological replicates. 5hmC containing DNA sequences were enriched from striatal total DNA and high-throughput sequencing resulted in a range of ∼20-70 million uniquely mapped reads from each biological replicate (**Supplemental Table 3**, methods). Similar to the hippocampal 5hmC data, sequence read density mapping showed no visible differences among the chromosomes, except depletion on chromosomes X, and an overall equal distribution of 5hmC levels in both genotypes and in both conditions on defined genomic structures and repetitive elements (**Supplemental Fig. 6**). To compare the long-lasting molecular effects of a gene by environment interaction on female mice in the striatum and hippocampus, we annotated the striatal HET-DhMRs to genes and found a significant overlap of differentially hydroxymethylated genes between both tissues (*N* = 389 genes; *P*-value < 0.05), including genes well known to play a role in neurodevelopmental disorders (*e.g., Nrxn1, Nlgn1,* and *Grip1/Ncoa2*; **Supplemental Fig. 7A**). Again, similar to the hippocampal findings, the gene ontologies of the striatal DhMR-associated gene contained a significant neuronal enrichment of terms (*e.g.,* synapse organization/transmission (glutamatergic) and negative regulation of axonogenesis; (*P*-value < 0.05 & FC > 1.5 for GO enrichment tests; Dataset 2). The DNA hydroxymethylation levels of several regions also were validated using an alternative method (**Supplemental Fig. 7B**; Methods). Together, these striatal 5hmC data provide a validation that 5hmC levels are stably disrupted following a GxE interaction in brain regions linked to stress response and social behavior.

Recently we reported that male *Cntnap2* knockout mice exhibit a genome-wide disruption of 5hmC in a significant number of orthologous genes and pathways common to human neurodevelopmental disorders^24^. Since the early-life stressed *Cntnap2^+/-^* female mice showed repetitive behaviors and altered sociability, similar to the homozygote phenotype, we compared the DhMR-associated genes between these two models (*e.g.,* ELS-HET and female homozygotes) and found a significant overlap of genes in the striatum (*N* = 443; *P*-value < 0.05; **Supplemental Fig. 7C**). In addition, the DhMR-associated genes from the homozygote striatum also had a significant overlap of gene when compared to the ELS-HET genes from the hippocampus (*N* = 356; *P*-value < 0.05; **Supplemental Fig. 7C**). Together, these data further validate the finding that 5hmC disruptions are linked to neurodevelopment-related disorders and support gene by environment hypotheses for complex disorders.

### Candidate functional DhMRs in the female GxE interaction model

To gain insight into the potential molecular mechanism(s) for long-lasting GxE interaction-induced DhMRs, we used RNA sequencing (RNAseq) to profile gene expression in the hippocampus of the same mice surveyed for 5hmC (see methods). Comparison of transcript levels in ELS-HET and control-HET mice revealed 524 differentially expressed genes, 162 upregulated and 363 downregulated (FDR *P*-value < 0.1; **Figure 3A**; Dataset 3). Separate gene ontologies of these 162 upregulated and 363 downregulated genes found terms related to metabolic-related processes (upregulated genes; **Figure 3B**; Dataset 4) and axonal development and histone modifications (downregulated genes; **Figure 3C**; Dataset 4; methods). DhMRs associated with differentially expressed genes represent candidate functional DhMRs that may have a direct role in gene regulation. An overlay of the HET-DhMR data with these HET-RNAseq data revealed that 72 differentially expressed genes harbor a HET-DhMR, which included neurodevelopmental disorder genes (*e.g., Reln*, *Dst,* and *Trio*)^41^, known stress-related genes (*e.g., Ncoa2*)^46^ and epigenetic genes (*Dnmt3a* and *Hdac4*; Dataset 3). While general relationships between 5hmC abundance and gene expression levels were not observed in these genes, there was a significant enrichment of DhMRs located in the introns of differentially expressed genes (*P*-value < 0.05). These results indicate that the GxE interaction induces long-lasting effects in the regulation of genes functioning in neuronal development and synaptic plasticity and supports an association between 5hmC and the altered social behavior observed in these adult mice.

**Figure 3:**
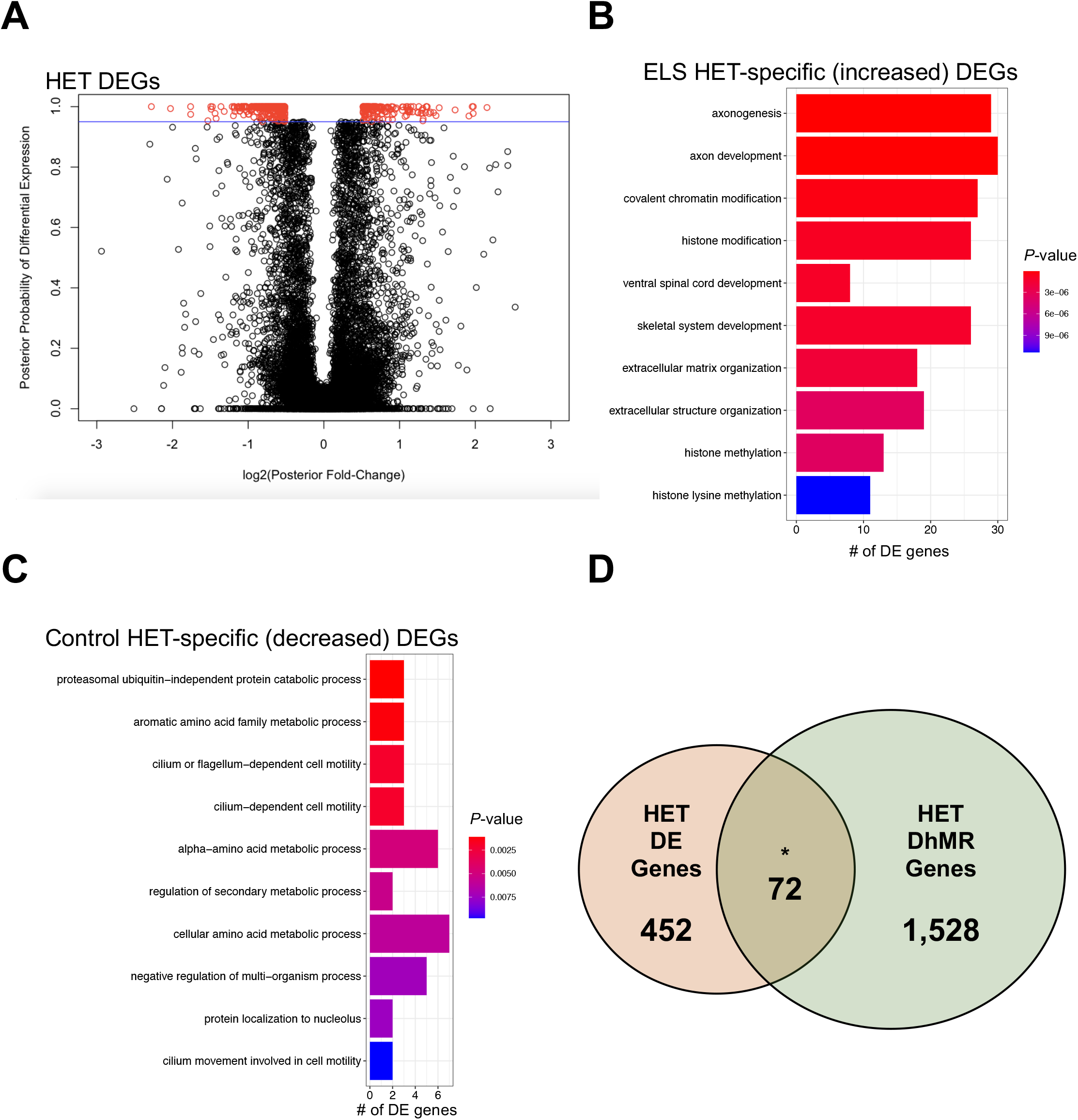
Characterization of differentially expressed genes (DEGs) between ELS-HET and Control-HET. (A) A modified volcano plot depicts the log2(posterior fold change; x-axis) versus the posterior probability of differential expression (y-axis). Open circles (red/black) represent each gene examined and differentially expressed genes (red) are shown above the significance line (blue, FDR *P*-value < 0.1). (B-C) A bar plot showing the top significantly over-represented ontological terms based on gene ontological analysis of ELS-HET-specific (B) differentially expressed genes and Control-HET-specific (C) differentially expressed genes. The number of DEGs linked to the ontological terms are displayed on the x-axis. Bar color is based on *P*-value. (C) A Venn diagram depicts the overlap of DhMR-associated genes (green circle) and differentially expressed genes (orange circle) from the ELS-HET versus Control-HET comparison. The asterisk (*) denotes a significant overlap (hypergeometric enrichment *P*-value < 0.05).

Comparison of transcript levels in ELS-WT and control-WT mice revealed only 5 genes that were altered by early-life stress, all downregulated (FDR *P*-value < 0.1; **Supplemental Fig. 8**; Dataset 3). An overlay of the WT-DhMR data with the WT-RNAseq data did not reveal any potentially functional DhMRs. These results suggest that WT-DhMRs represent stable molecular alterations that may have affected gene expression at earlier developmental time-points, suggesting that the ELS-WT mice may have earlier phenotypes related to the early-life stress.

### The GxE interaction alters transcription factor binding in DhMRs

Syndromic forms of neurodevelopmental disorders can involve the disruption of transcription factor function^47, 48^; thus, we investigated the potential for the HET-DhMRs to alter DNA-binding of transcription factors by testing for enrichments of known transcription factor sequence motifs among the DhMRs (Methods). Several transcription factor sequence motifs were significantly enriched in the hippocampal HET-DhMRs. Many of the transcription factors that bind these motifs have links to neurodevelopmental behaviors and disorders, such as CLOCK, NPAS2, HIF-1A/B, and USF1 (**Table 2**; Dataset 5)^49,50,51,52^. Together, these findings suggest that 5hmC influences transcription factor binding, which may explain the observed correlated disruptions in gene expression. An initial test of this hypothesis examined transcription factor sequence motif enrichments among only the potentially functional DhMRs (*i.e.,* genes with correlated disruptions in 5hmC and expression; *N* = 72; Methods). Five transcription factors sequence motifs were significantly enriched among the potentially functional HET-DhMRs (bHLHE40, c-MYC, CLOCK, HIF-1b, and USF1). The CLOCK transcription factor binds E-box enhancer elements, its consensus binding sequence contains a CpG dinucleotide (CACGTG) that can be hydroxymethylated, and its expression/dysregulation is associated with autism^53, 54^. RNA sequence data and Western blot analysis revealed that *Clock* is not differentially expressed at the transcript and protein levels due genotype and/or condition (*i.e.,* ELS-HET, control-HET, ELS-WT, and control-WT; (**Figure 4A**; Dataset 3). To determine the effect of this GxE interaction on the genome-wide distribution of CLOCK binding, we employed chromatin affinity purification coupled with high-throughput sequencing technology (ChIP-seq). Age-matched ELS-HET and control HET, as well as ELS-WT and control-WT female offspring (*N* = 3/group) were sacrificed at 3 months of age as independent biological replicates. DNA sequences containing CLOCK binding sites were enriched from the hippocampus total DNA and high-throughput sequencing resulted in ∼10 million uniquely mapped, non-duplicated, reads from each biological replicate (methods). A differential analysis of these data filtered to only DhMRs containing a putative CLOCK binding site (*N* = 577; CACGTG) identified 19 DhMRs with differential CLOCK binding (**Figure 4B**; Dataset 6).

**Figure 4:**
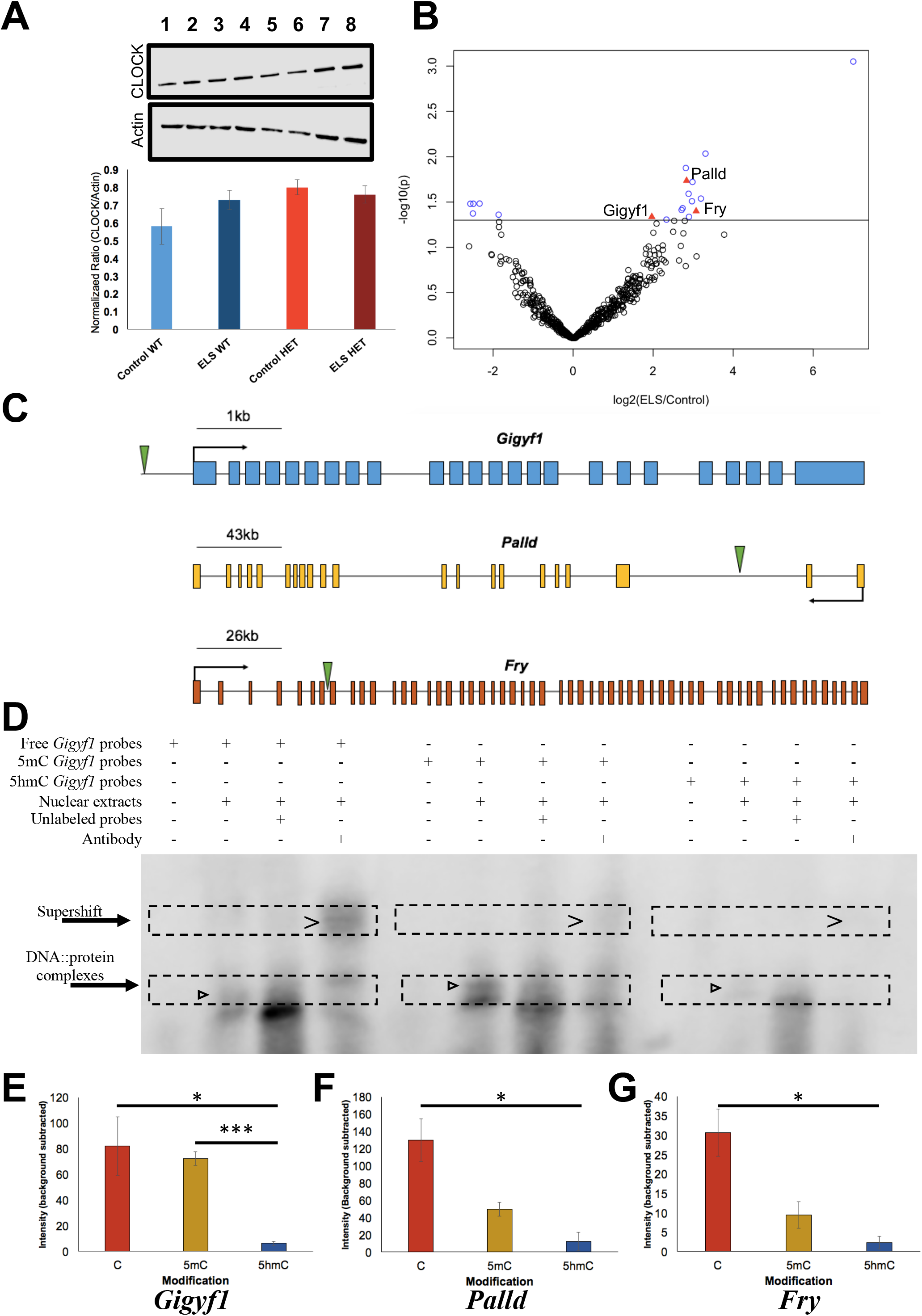
Functional assays reveals that 5hmC represses CLOCK binding. (A) Western blot analysis identified no gross differences in CLOCK protein expression between Control-WT (lanes 1 and 2), ELS-WT (lanes 3 and 4), Control-HET (lanes 5 and 6), or ELS-HET (lanes 7 and 8) (top panel) or in beta-actin (middle panel), used as a loading control in all lanes. A bar plot (bottom panel) depicts the quantified levels of CLOCK proteins normalized to beta-actin from each respective lane. Error bars show the standard error of the mean (*N* =2). (B) A volcano plot displays the –log2(fold-change) of ELS-HET compared to Control-HET (x-axis) and the significance (y-axis) from differential analysis of ChIP-seq data. Open circles (blue/black) and red triangles represent differentially hydroxymethylated regions (DhMRs) containing a canonical CLOCK binding site. DhMRs with significant differential binding of CLOCK (blue open circles and red triangles) are located above the significance line (*P*-value <0.05). Red triangle DhMRs also exhibit differential gene expression (*Gigyf1*, *Palld*, and *Fry*). (C) Gene schematics of *Gigyf1* (blue)*, Palld* (yellow), and *Fry* (orange) are shown and indicate the location of the DhMRs containing the differential CLOCK binding (green triangle). (D) Electromobility shift assays with hippocampal lysates. Lysate shift (lanes 2, 6, and 10), cold competition (lanes 3, 7, and 11), and supershift (lanes 4, 8, and 12) of DNA probes corresponding to a CLOCK E-box motif in the DhMR of *Gigyf1* that exhibited differential CLOCK binding. Biotinylated DNA probes contained either an unmodified E-box motif (lanes 1-4), a methylated E-box motif (lanes 5-8), or a hydroxymethylated E-box motif (lanes 9-12). Regions containing bands of interest are highlighted (dashed boxes) with DNA::protein complexes that are abrogated with cold competition are indicated (black arrowheads) and supershift complexes are shown (carrot). (E-G) Quantifications of biotinylated DNA probe intensities. The E-box modifications (x-axis) and mean biotinylated DNA probe intensity (y-axis) are depicted +/- SEM for *Gigyf1* (E), *Palld* (F), and *Fry* (G) probes. The asterisk (*) denotes a significant decrease in CLOCK binding box (* *P*-value <0.05; *** *P*-value <0.001).

**Table 2:**
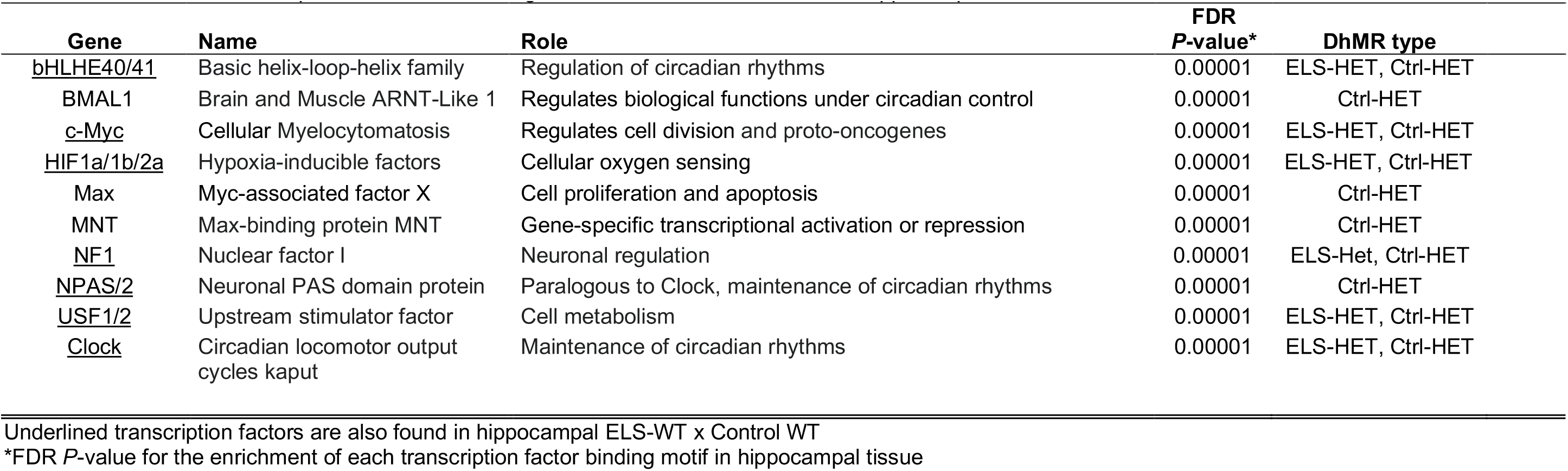
A subset of the transcription factors with binding motifs associated with DhMRs in hippocampus ELS-HET x Control-HET

Three of these DhMRs were associated with genes that also were differentially expressed as a result of the GxE interaction (*Gigyf1*, *Palld*, and *Fry*). The DhMR containing CLOCK binding sequences reside near the promoters of each of these genes (**Figure 4C**) and all three genes have links to autism.^55,56,57,58^ Together, these data suggest that differential hydroxymethylation levels may mediate CLOCK binding, supporting a mechanism for 5hmC in the regulation of gene expression of genes associated with the observed phenotype.

### 5hmC alters CLOCK binding

To formally test whether 5hmC can alter CLOCK binding, hippocampal nuclear lysates were incubated with oligonucleotide probes containing sequences from the differentially bound CLOCK sites in the DhMRs associated with *Gigyf1*, *Palld*, and *Fry* genes. The CpG dinucleotide in the E-box motif of CLOCK located in each probe contained either cytosine, 5-methylcytosine, or 5-hydroxymethylcytosine (Methods). Electromobility shift assays using unmodified cytosine probes from each of these genes revealed successful binding of CLOCK and BMAL1 from hippocampal nuclear lysates (**Figure 4D-G**; **Supplemental Fig. 9**). These DNA::protein complexes were abrogated in cold completion assays and super-shifted with the addition of antibodies against CLOCK (**Figure 4D**, left panel; Supplemental Fig. 9). In contrast, hydroxymethylated probes significantly reduced the binding of CLOCK (**Figure 4D-G**; **Supplemental Fig. 9**), indicating that 5hmC prevents CLOCK from binding to the E-box motifs located near the promoters of the differentially expressed genes *Gigyf1*, *Palld*, and *Fry*.

Together, these findings suggest a working model for 5hmC regulation of developmentally important genes that when altered can result in social-related disorders. In the example presented here, the GxE interaction results in a loss of 5hmC near the promoters of *Gigyf1*, *Palld*, and *Fry* that subsequently improves the binding of CLOCK to act as a repressor of *Gigyf1*, *Palld*, and *Fry* expression (**Figure 5)**. Thus, *Gigyf1*, *Palld*, and *Fry* are top candidate genes contributing to the adult sex-specific phenotype found in female *Cntnap2* heterozygous mice exposed to early-life stress.

**Figure 5:**
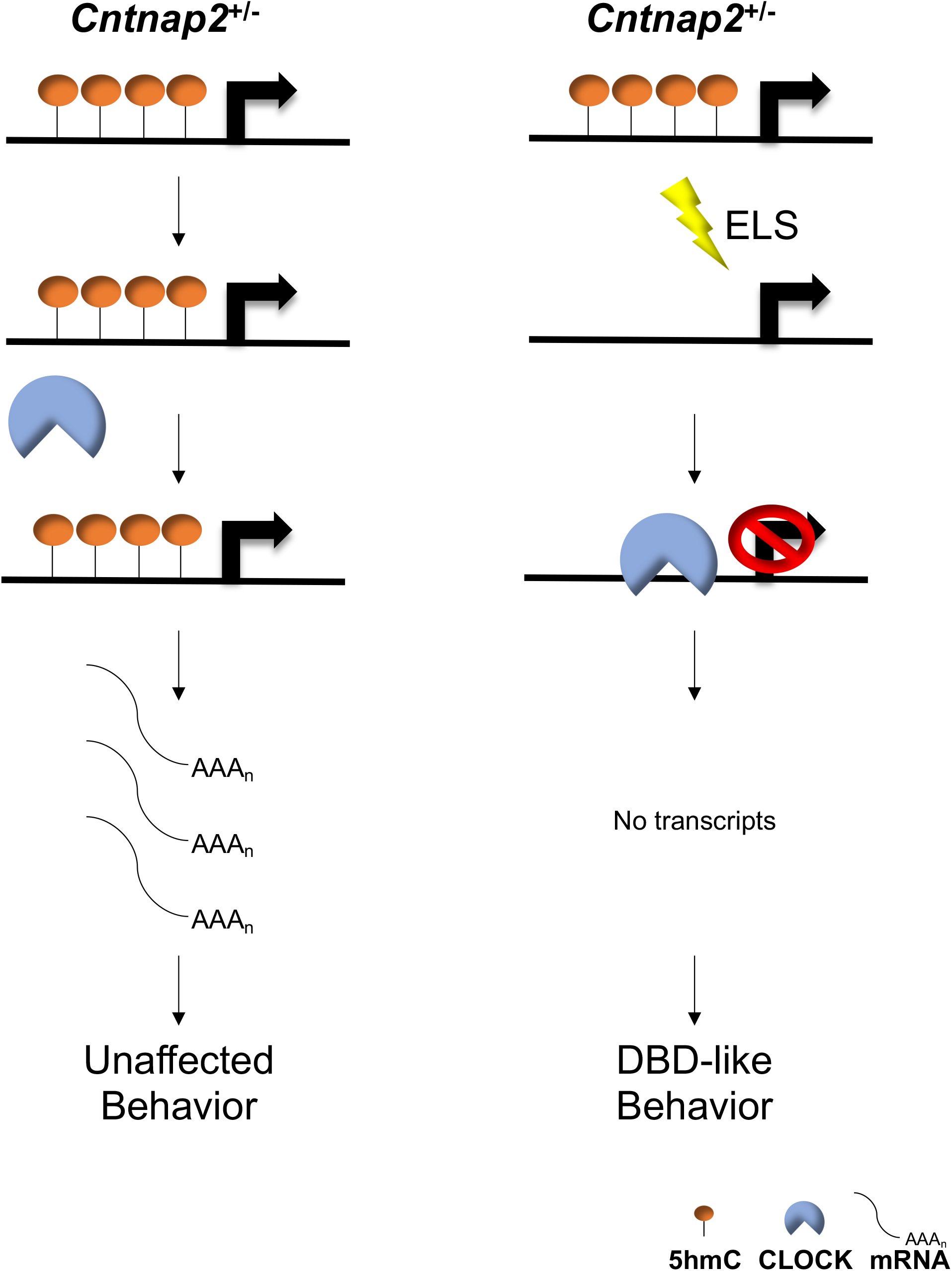
A working model of a functional role for 5hmC in developmental brain disorder-like behaviors. Control *Cntnap2^+/-^* mice (left panel) 5hmC levels (orange lollipops) remain abundant and prevent CLOCK (blue semicircle) from binding to genes implicated in developmental brain disorders (*e.g., Gigyf1*), which results in adequate gene expression levels (black curved lines) and unaffected behaviors. *Cntnap2^+/-^* mice exposed to early life stress (ELS; lightning bolt; right panel) exhibit a loss of 5hmC abundance in genes implicated in developmental brain disorders (*e.g., Gigyf1*). This reduction in 5hmC levels allows CLOCK to bind to DNA and repress transcription, which results in developmental brain disorder-like phenotypes.

## Discussion

Here we provide evidence of a functional role for 5hmC in a gene (partial loss of *Cntnap2*) by environment (early-life stress; GxE) interaction model that results in sex-specific alterations of neurodevelopmental behaviors. These 5hmC disruptions were found in the orthologs of a significant number of genes implicated in human developmental brain disorders: significantly overlapping with those found in both male and female *Cntnap2* homozygous mouse mutants^59^. Integration of epigenetic and gene expression data from the same mice revealed relations with genes (*e.g., Nrxn1*, *Reln*, *Grip1*, *Dlg2* and *Sdk1*) known to contribute to the behavioral deficits of neurodevelopmental disorders. Finally, we focused on an enrichment of transcription factors sequence motifs among the potentially functional ELS-HET DhMRs and found that the loss of 5hmC in these regions resulted in the loss of CLOCK binding. Together these data demonstrate a molecular mechanism for 5hmC in the regulation of developmentally important genes that when altered can result in social-related disorders. A GxE model and 5hmC disruptions may represent a common etiology for developmental brain disorders in humans.

Having data from both striatal and hippocampal tissues provide a unique opportunity to examine the combined and differential contributions of these tissues to neurodevelopment-related behaviors, which both supported that the epigenome plays a role in the long lasting effects of early-life environmental exposures on brain and behavior. For example, both tissues carried GxE DhMRs in orthologous genes common to human neurodevelopmental disorders, such as nuclear receptor genes, glucocorticoid receptor-interaction protein 1 (*Grip1* also called *Ncoa2*)^46^, as well as several genes associated with neurotransmission and synaptic plasticity (*Nrxn1*, *Nlgn1*, *Dlg2*, and *Negr1*)^41, 60^. The hippocampus-specific GxE DhMRs also included genes related to human neurodevelopmental disorders, including *Homer1*, which forms high-order complexes with *Shank1* that are necessary for the structural and functional integrity of dendritic spines^61^. In addition, the hippocampus-specific GxE DhMRs contained links to circadian function (*e.g., Clock*, *Homer1*, and *Shank2*); the prenatal stress administered here is sleep disruptive, supporting a hypothesis that circadian rhythms contribute to the pathogenesis of neurodevelopmental disorders^62^. Finally, investigation of the hippocampal-specific GxE DhMRs with correlated disruptions in gene expression revealed links to the observed adult behavioral deficits, including specific links to the limbic system and synaptic plasticity (*e.g., Dst, Reln, Lpp, Trio*, *Notch1 Col4a1*, and *Utrn*)^63,64,65,66^. Together, these data suggest insights into the molecular pathogenesis of GxE interactions that result in neurobehavioral alterations.

Importantly, while early-life stress alone disruptions in 5hmC (WT-DhMRs) were largely unique from the GxE DhMRs (HET-DhMRs), WT-DhMR-associated genes also included previous links to neurodevelopmental disorders (*e.g., Auts2*, *Cacna1c* and *Setd5*)^41^. Although 5hmC disruption on these genes was not correlated with altered transcript levels and were not sufficient to cause a behavioral phenotype, many of them have been implicated in disorders with social deficits^41^, suggesting that early-life stressed WT mice have uncharacterized deficits and may be susceptible to additional environmental stressors that trigger later life brain-related disorders.

Mechanistically, it is intriguing that the binding motifs of several transcription factors with known roles in neuronal development were found in the GxE DhMRs. These findings included binding motifs for transcription factors best known for their roles in circadian rhythm, immune response, learning and memory, and hypoxia. For example, CLOCK, BMAL, BHLHE40, and BHLHE41 all regulate the circadian clock; in addition, they are also linked to neurodevelopmental disorders, such as major depressive disorders, bipolar disorders, and autism spectrum disorders^50,51,52, 67^; again, consistent with the prenatal stress paradigm used here being sleep disruptive. CLOCK functions as a pioneer-like transcription factor^68^, while also displaying histone acetyltransferase activity^69^, corroborating CLOCK as a regulator of chromatin accessibility and contributor to the epigenetic framework in cells. While CLOCK generally acts as a transcriptional activator, it also exhibits repressive activity that is dependent on the assembly of larger protein complexes^70,71,72^. Paired box protein 5 (*Pax5*) and aryl hydrocarbon receptor (*Ahr*) have defined roles in immune response and also are connected to autism spectrum disorders and major depressive disorders^67, 73, 74^. The stress paradigm used in this study increases placental inflammation^75^; indeed, immune activation history in the mother is associated to increased symptom severity in children with mental illness^76^. It is notable that the GxE interaction resulted in distinct differences between ELS-HET and ELS-WT enriched transcription factor sequence motifs, perhaps suggesting an interaction between *CNTNAP2* and 5hmC in transcription factor binding and function.

Focusing more specifically on significant enrichments of transcription factor sequence motifs in potentially functional DhMRs, especially enriched motifs located in the promoter regions, revealed known and novel genes contributing to the adult phenotype. *Gigyf1*, *Palld*, and *Fry* were the examples reported here with differential CLOCK binding and links to autism.^55,56,57,58^ In addition, *Catenin Alpha 2* (*CTNNA2*) is an example of a gene that is differentially expressed and contains a DhMR located in its promoter harboring a different enriched transcription factor sequence motif (USF1). *Ctnna2* encodes a protein that regulates morphological plasticity of synapses and its loss in neurons leads to defects in neurite stability and migration, and cortical dysplasia^77^. Indeed, cortical dysplasia is a feature in humans and mice lacking expression of *Cntnap2*, and mice exhibit defects in neuronal migration of cortical projection neurons and a reduction in the number of striatal GABAergic interneurons^27, 28^. It will be important to characterize the role of 5hmC on the binding of other transcription factors (*e.g.,* USF1) in the promoters of genes containing potentially functional DhMRs. While enriched motifs in DhMRs located in promoter regions of differentially expressed genes represent top candidate genes directly contributing to the adult phenotype, enriched motifs in DhMR-associated genes that are not differentially expressed in the adult may identify genes contributing to the origins of the adult behavioral deficits. Further studies at earlier developmental time-points are needed to determine the role of these DhMRs in the development of the observed adult behavioral deficits.

Exposure to stress during different gestational periods has unique effects on epigenetic programming of the developing embryo. During early gestation, early-life stress-induced epigenetic changes in neuronal precursor cells are predominantly mediated by hormonal mechanisms, which in part can be transmitted via placental pathways^78^. Stress during this time period predominately affects behavioral development in males^30, 79^. However, in late gestational stages, the mechanism underlying early-life stress-induced epigenetic changes become more complex involving the activation of the excitatory and inhibitory endocrine modulatory systems, which may interfere with stress-induced synaptic reorganization^78^. Stress during this time period predominately affects behavioral development in females^30, 79^; thus, there is a sex-specific effect on behavior depending on the gestational timing of the stress exposure. Consistent with these reports, late gestational stress in this present study results in long-lasting behavioral and molecular effects in females on genes known to be associated with synaptic plasticity and mental disorders. The repetitive behaviors and social deficits render the ELS-HET mice an attractive model for other psychiatric disorders in humans. Since both insults (*i.e.,* genetic and environment) are required for the expression of an altered behavioral phenotype, this model (*i.e.,* a two-hit model) directs studies to consider environmentally sensitive molecular mechanisms contributing to stress-induced behavioral outcomes. Moreover, it supports the study of other “multi-hit” hypotheses in the origins of mental illness to identify elusive factors that could contribute to the complexity of psychiatric disorders in humans. While there is precedence for multi-hit models disrupting brain development^80,81,82,83^, it will be important to examine the generalizability of these findings using other developmentally important genes in the brain that also may contribute to the vulnerability/resilience toward mental illness.

Importantly, *CNTNAP2* has been implicated in numerous neurodevelopmental disorders and heterozygous disruptions have been found in humans with intellectual disability (ID), seizures, and signs of autism spectrum disorders (*e.g.,* repetitive behaviors, and social deficits). However, heterozygous losses of *CNTNAP2* also have been inherited from healthy parents, suggesting that heterozygous disruptions of *CNTNAP2* by themselves are not sufficient to elicit cellular or organismal phenotypes. Aside from striatum and hippocampus, high levels of *CNTNAP2* expression have been found in the olfactory bulb, ventricular zone, and thalamus^28, 84^, warranting future examinations of 5hmC in other brain-related behaviors that are linked to these additional brain regions. *CNTNAP2* also has an organizing function to assemble neurons into neural circuits^85^, suggesting future studies should examine the electrophysiology of the excitatory and inhibitory synaptic transmissions in the ELS-HET mice. Finally, early-life stress may have revealed a molecular connection between 5hmC and *Cntnap2* involving the binding of a transcription-enhancing transcription factor *TCF4,* which can transactivate the *CNTNAP2* and *NRXN1* promoters. Heterozygous deletions of *TCF4* causes Pitt-Hopkins syndrome: a syndrome that also is linked to the loss of either *CNTNAP2* or *NRXN1* function^86,87,88^. Thus, *TCF4* may modulate the expression of *CNTNAP2* and *NRXN1* in the regulatory network involved in Pitt-Hopkins syndrome^89^. Interestingly, we find several DhMRs associated with *Tcf4* and *Nrxn1* following GxE interaction, suggesting that the regulatory network involved in Pitt-Hopkins syndrome may be similarly disrupted in the ELS-HET mice. Together, these findings potentially provide novel insight into the molecular connection between 5hmC, *Cntnap2*, *Tcf4*, and *Nrxn1* in neurodevelopmental disorders, such as Pitt-Hopkins syndrome.

Molecular factors influencing vulnerability and resilience to environmental stress clearly involve environmentally sensitive epigenetic mechanisms such as DNA methylation. Thus, perturbations of gene by environment interactions may underlie common pathways for many mood-related disorders in humans. The identification and characterization of a potentially modifiable substrate (*e.g.,* 5hmC) contributing to gene by environment interaction-induced sex-specific neurologic behaviors is significant and comes at a time when there is great interest to harness the diagnostic and therapeutic power of these substrates toward healthy outcomes.

## Methods

### Mice and genotyping

All experiments were approved by the University of Wisconsin – Madison Institutional Animal Care and Use Committee (M02529). Heterozygous male *Cntnap2^+/-^* mice were purchased from the Jackson laboratories (Bar Harbor, ME) and maintained on C57BL/6J background, as previously reported^24^. *Cntnap2^+/-^* mutants were genotyped using the following primers: Mutant Rev: CGCTTCCTCGTGCTTTACGGTAT, Common: CTGCCAGCCCAGAACTGG, WT Rev 1: GCCTGCTCTCAGAGACATCA. PCR amplification was performed with one cycle of 95°C for 5 min and 31 cycles of 95°C for 30 sec, 56°C for 30 s, 68°C for 30 s, followed by 68°C for 10 min. The mutant allele was obtained with a 350-bp and wild-type (WT) allele with a 197-bp PCR products.

### Breeding scheme and prenatal stress paradigm

To minimize the stress of animal handling, all of the following were conducted by a single researcher: animal colony maintenance; breeding, prenatal stress; and behavioral tests. For breeding, a three month old *Cntnap2*^+/-^ male mouse was placed together with a virgin 3 month old WT C57/Bl6J female mouse at 6 pm (one hour prior to lights off); every morning before 9 am (two hours after lights on) female mice were checked for vaginal copulation plug and separated from male. Presence of a copulation plug denoted day 1 of gestation and the pregnant female was individually housed and given a cotton nestlet. At day 12 of gestation (E12), pregnant females were randomly assigned to either a variable stress or non-stressed control group. Pregnant mice assigned to the variable stress group experienced a different daily stressor on each of the seven days during late pregnancy (E12 to E18; the timing of which was chosen because it overlaps with the onset of *Cntnap2* expression (E14)). These variable stressors included: 36 hours of constant light, 15 min of fox odor (Cat# W332518) beginning 2 hours after lights on, overnight exposure to novel objects (8 marbles), 5 minutes of restraint stress (beginning 2 hours after lights on), overnight novel noise (white noise, nature sound-sleep machine®, Brookstone), 12 cage changes during light period, and overnight saturated bedding (700 mL, 23°C water)^31, 32^. These mild stressors were previously published and were selected because they do not include pain or directly influence maternal food intake or weight gain.

Importantly, both WT and HET mice were taken from the same litter, providing an ideal internal control for our findings and negates the need for cross-fostering of the prenatally stressed pups to non-stressed moms to find effects of early-life exposure to stress. Litter sizes of less than 5 and more than 8 pups were removed from the experiment. Offspring were ear-tagged at postnatal day (P) 12 and left undisturbed until weaning day (P18), at which time the mice were group housed with same sex. Female offspring were left intact, and were not cycled^32^. Finally, to minimize the effect of parent-to-offspring interaction per litter, a maximum of 3 pups/litter were randomly selected for the behavioral or molecular experiments (total of *N* = 16 litters for both experimental and validation cohorts). Importantly, the mice used for behavioral and molecular analysis were left undisturbed until behavioral testing or sacrifice at 3 months of age.

### Behavioral Tests

Early-life stressed and non-early-life stressed *Cntnap2^+/-^* and WT mice (both sexes, 12-17 weeks of age) were submitted to tests of anxiety, depression, and sociability. All testing was performed during a period of 4 hours in dark period (beginning 2 hours after lights off). With one-week between tests, the same group of mice were subjected to the following sequence of behavioral tests: open field, light/dark box, elevated plus maze, forced swim test, 3-chamber social test. For the validation of the repetitive behavior and social deficits the same group of mice were subjected to the following sequence of behavioral tests, with one-week between tests: ten-minute observational test, 3-chamber social test, and Ten-minute reciprocal social interaction test. An experimenter who was blind to the animal’s group and genotype scored video recordings of each test.

#### Light/Dark Box Test

The light/dark box test was performed in a rectangular box divided in two compartments (light and dark). The walls of the light and dark compartments were constructed of Plexiglass (same areas 319.3 cm^2^). A removable dark Plexiglass partition was used to divide the box into light and dark sides. Each animal was placed into the light side of the box, facing away from the dark side and allowed to explore both chambers of the apparatus for 10 min. The time spent in the light side was scored for each mouse.

#### Open Field Test

The open field apparatus consisted of a circular arena (104 cm diameter). A marker was used to inscribe a smaller circle 25 cm from the walls. Each mouse was placed in the inner circle of the apparatus and allowed to explore for 10 min. The time spent in the center and numbers of entries to inner cycle were scored for each mouse.

#### Elevated Plus Maze

The elevated plus maze consisted of two open and two closed arms, elevated 52 cm above the floor, with each arm projecting 50 cm from the center (a 10 × 10 cm area). Each mouse was individually placed in the center area, facing the open arm, and allowed to explore for 7 min. The measurements scored for each mouse were as follows: time spent in closed arms, time spent in open arms, and time spent in center.

#### Forced Swim Test

The mice were individually placed into a glass cylinder (14-cm internal diameter, 38 cm high) filled with water (28-cm deep, 25–26 °C) for 6 min. During the last 4 min of the test, the time spent floating was scored for each mouse. Floating was defined as immobility or minimal movements necessary to maintain the head above the water.

#### 3-Chamber Social Test

The social interaction test was performed as previously described^90^. Briefly, an experimental mouse was placed in the center third of a Plexiglass box (77.5 x 44.4 cm, divided into three equal chambers) for 10 minutes. Next the experimental mouse was allowed to explore all three interconnected chambers for 10 minutes. Finally, an empty wire cup was placed in one of the end chambers and an identical wire cup containing an unfamiliar mouse of the same sex was placed in the chamber at the opposite end; the experimental mouse was allowed to explore all three interconnected chambers for 10 minutes.

Time spent in each chamber and spent sniffing each cup as well as number of entries to each chamber was measured for each mouse.

#### Ten-minute reciprocal social interaction test

Mice were placed in a cage (previously habituated to it) with an unfamiliar mouse matched for treatment, genotype, and sex for 10 minutes. The time mice were engaged in repetitive behaviors (grooming and digging) was measured.

#### Ten-minute observational test

Mice were individually placed in a cage and the time engaged in repetitive behaviors (grooming and digging) was measured.

### Statistics for behavioral tests

Data is reported as mean ± S.E.M. Homogeneity of variance was assessed using the Brown-Forsythe and the normal distribution of the data was tested by the Shapiro-Wilk test. Two-way ANOVA also was used to detect differences between genotype (WT x HET) and treatment (Control x ELS) for time spent in the inner circle and number of entries into that circle (Open field test); time spent in closed arm, time spent in open arm, and time spent in center (Elevated plus maze); time spent in light side (Light/Dark box test) and time spent floating (Forced swim test). Two-way ANOVA was used to detect differences between treatment (Control x ELS) for time spent in the chamber (3-chamber social test) and interacting with the novel mouse and time spent in the chamber with the empty cup. One-way ANOVA was used to detect differences between groups for time spent engaged in repetitive behaviors (grooming and digging). Post hoc analysis was performed using a Bonferroni correction. Statistics were performed using SigmaPlot version 14.

### DNA and RNA extractions

DNA methylation and gene expression was obtained from early-life stressed and non-early-life stressed three-month old female *Cntnap2^+/-^* and WT mice (*N* = 3 per group). Mice were sacrificed (2 hours after lights on) and whole brains were extracted without perfusion and immediately flash frozen in 2-methylbutane and dry ice. Striatum tissue was excised by micropunch (1.53 to -0.95 mm posterior to bregma), while whole hippocampal tissue was excised by morphology. Approximately 30 milligrams of tissue was homogenized with glass beads (Sigma) and DNA and RNA were extracted using AllPrep DNA/RNA mini kit (Qiagen).

### 5hmC Enrichment of Genomic DNA

Chemical labeling-based 5hmC enrichment was described previously^15, 24^. Briefly, a total of 10ug of striatum DNA was sonicated to 300 bp and incubated for 1 hour at 37°C in the following labeling reaction: 1.5 ul of N3-UDPG (2mM); 1.5ulβ of -GT (60uM); and 3ul of 10X β-GT buffer, in a total of 30ul. Biotin was added and the reaction was incubated at 37°C for 2 hours prior to capture on streptavidin-coupled dynabeads (Invitrogen, 65001). Enriched DNA was released from the beads during a 2 hour incubation at room temperature with 100mM DTT (Invitrogen, 15508013), which was removed using a Bio-Rad column (Bio-Rad, 732-6227). Capture efficiency was approximately 5-7% for each sample.

### Library Preparation and high-throughput sequencing

5hmC-enriched libraries were generated using the NEBNext ChIP-Seq Library Prep Reagent Set for Illumina sequencing, according to the manufacturer’s protocol. Briefly, the 5hmC-enriched DNA fragments were purified after the adapter ligation step using AMPure XP beads (Agencourt A63880). An Agilent 2100 BioAnalyzer was used to quantify the amplified library DNA and 20-pM of diluted libraries were used for sequencing. 50-cycle single-end (striatum) or paired-end (hippocampus) sequencing was performed by Beckman Coulter Genomics or the University of Wisconsin Biotechnology Center, respectively. Paired-end sequencing for hippocampal tissue was chosen for reasons unrelated to this project. Image processing and sequence extraction were done using the standard Illumina Pipeline.

### Analysis of 5hmC data: sequence alignment, fragment length estimation and peak identification

We mapped the reads to mouse NCBI37v1/ mm9 reference genome using Bowtie 0.12.7^91^, allowing for no more than two mismatches throughout the entire read and only keeping the uniquely mapped reads. For striatal data: the Model-based Analysis of ChIP-Seq 2 (MACS2) algorithm v2.1.2 ^92^ was used to estimate fragment size, call peaks, and identify peak summits from aligned single-end reads using the following parameters: single-end format, effective genome size of 1.87e9, band width of 300bp, an FDR cutoff of 0.01, auto pair model process enabled, local bias computed in a surrounding 1kb window, and a maximum of one duplicate fragment to avoid PCR bias. For hippocampal data: MACS2 also was used, employing the same parameters except in the paired-end mode, and using an FDR cutoff of 0.05, which could be relaxed (compared to striatal data) due to having paired-end sequence data, which increases the signal in the data. Summits from both striatal and hippocampal data were extracted for each peak for each sample and extended +/-500bp for downstream analysis. We defined the peaks for each group as follows using the peaks from all three subjects: peaks were merged from all subjects in a group and group peaks were identified if individual peaks overlapped between a minimum of two subjects in the group.

### Identification of differentially hydroxymethylated regions (DhMRs)

For each genotype, the stress and control groups were pooled and merged to form the candidate differential regions within each genotype. The Bioconductor package edgeR was used to test whether a difference of read counts exists between the two groups in each candidate region (*P*-value<0.05)^93^. The types of DhMRs (specific to each genotype/condition) were determined by the average log fold change in a normalized read count (logFC) between the early-life stressed mice and controls.

### Annotation of sequence reads, peaks, and DhMRs

All the reads were extended to their fragment lengths estimated by MACS2. We extracted genomic features and their associated gene symbols from GENCODE M1 and R Bioconductor package org.Mm.eg.db^94^, and downloaded the repetitive elements from UCSC genome browser (https://genome.ucsc.edu/)95. We used Binomial tests to determine the significance in read density and DhMR distribution differences over the chromosomes, genomic features and repetitive elements. When conducting the binomial test for DhMRs, the background proportions were calculated from all the peaks in the two groups of animals in each comparison. We used ngsplot to draw the profile plots of the genotype/condition 5hmC enrichments and other peaks^96^.

### Simultaneous targeted methylation sequencing for molecular validation

Molecular validation of DhMR data was performed as previously described^97^. PCR-amplicons were sequenced on an Illumina miSeq following standard protocols. Illumina adapter sequences were removed from paired-end FASTQ files using Trim Galore v0.4.4. Alignment of trimmed reads was performed using Bismark v0.19.0, coupled with Bowtie2 v2.2.0^91, 98^, using a seed mismatch parameter of one base pair, a maximum insertion length of 1000bp, with all other parameters set to default. Mapping efficiency ranged from ∼78-85%. Read coverage and methylation calling was extracted using bismark_methylation_extractor with the following parameters: paired-end files used, no overlapping reads reported, with methylation output reported as a sorted bedGraph file^97^. Coverage files were imported to R environment and *methylKit* was used to determine differentially hydroxymethylated loci (DhMLs)^99^. A *P*-value threshold of 0.05 was used to determine significance.

### RNA sequencing

Approximately 100 ng of total RNA was used for sequence library construction following instructions of NuGen mRNA sample prep kit (cat# 0348). In brief, total RNA was copied into first strand cDNA using reverse transcriptase and random primers. This was followed by second strand cDNA synthesis using DNA Polymerase I and RNaseH. These cDNA fragments went through an end repair process, the addition of a single ‘A’ base, and ligation of adapters. These products were gel purified and enriched with PCR to create the final cDNA libraries. The library constructs were run on the bioanalyzer to verify the size and concentration before sequencing on the Illumina HiSeq2500 machine where 100-cycle single-end sequencing was performed by the University of Wisconsin Biotechnology Center. In total, three libraries (the same mice and combinations that were used to generate the 5hmC data) were sequenced for each experimental condition.

### Analysis of RNA-sequence data

Read alignment and calculation of transcript expression levels: we used the mm9 assembly as our reference genome and the GENCODE M1 (NCBIM37) as our gene annotation library (the Y chromosome was deleted for the alignment of female samples). We ran RSEM to calculate the expression at both gene-level and isoform-level, where RSEM automates the alignment of reads to reference transcripts using the Bowtie aligner to assign fractional counts for multi-mapping reads^100^. Bowtie was set to report all valid hits with up to 2 mismatches allowed, and to suppress all the alignments if a read has more than 200 hits. Differentially expressed (DE) genes/transcripts detection: EBSeq was used to detect the DE genes and transcripts in each of the pairwise comparisons of ELS-WT vs control WT, and ELS-HET vs control HET. We tested on both the gene level and the isoform level with a maximum number of differentially expressed groups of isoforms to be three for each gene. Genes or transcripts with more than 75% values < 10 (*i.e.,* 4 out of 6 samples in each contrast) are filtered to ensure better model fitting, and the normalization factors are the median count for each gene/transcript. We applied a FDR control to the testing and picked out genes/transcripts with FDR < 0.1.

### Enrichment tests of genes and GO analysis

DhMRs were annotated to all genes within 10k. To test for the enrichment of known neurodevelopmental genes among the DhMR-associated genes, we used a chi-square test to compare the DhMR-associated genes to a list of orthologs of well-documented human developmental brain disorder genes (*N* = 232)^41^. Bioconductor package clusterProfiler was used to test GO Biological Process (BP) term enrichment with *P*-value cutoff of 0.05 and an enrichment fold-change > 1.5^101^. For the GO enrichment analysis of DhMR associated genes, the gene universe consisted of all the genes associated with 5hmC peaks in both the early-life stressed and the control mice. For the GO analysis of DE genes, the gene universe is all genes that survived the filtering of EBSeq. To test for an enrichment of neuronal related GO BP terms among the DhMR-associated GO BP terms, we used a chi-square test and a previously published list of neuronal related GO BP terms (*N* = 3,046)^102^.

### Sequence motif analysis

For motif discovery analysis, the Hypergeometric Optimization of Motif EnRichment (HOMER) suite of tools was utilized^103^. DNA sequences corresponding to DhMR coordinates were obtained from the mm9 genome and compared against background sequences, using the given size of the region. Enriched known motifs of vertebrate transcription factors (*N* = 428) were determined using binomial testing and an FDR cutoff of 0.05.

### Western blot

Mouse hippocampal nuclear extracts were isolated from adult female mice (P90) using a nuclear and cytoplasmic extraction kit per the manufacturer’s instructions (Thermo Scientific 78833). Forty micrograms of total protein from hippocampal nuclear lysates was boiled for 5-minutes to dissociate complexes before separation in a 4-20% gradient gel (Bio-Rad 4561096) for 1-hour at 150V in 1x Tris/Glycine/SDS (Bio-Rad 1610732). Following separation of proteins, gels were transferred to nylon membrane in 1x Tris/Glycine for 1-hour on ice at 100V. Following transfer, membranes were washed in 1X TBST and blocked in 5% milk/TBST for 1-hour at room temperature. Blocked membranes were incubated with primary antibodies for 12-hours at 4°C. Primary antibodies included: 1:200 dilution anti-CLOCK (Abcam ab3517) and 1:4000 dilution anti-Actin (Abcam ab8226). Excess primary antibody was removed and membranes were washed in 1x TBST for 1-hour at room temperature. Washed membranes were incubated with secondary antibody (Abcam ab97051) in 5% milk/TBST at room temperature.

Membranes were washed at room temperature 1-hour with 1x TBST. Chemilumiescence was achieved with the Pico Chemiluminescence Kit (Thermo Fisher 34579) following manufacturer’s instructions. Visualization was achieved utilizing an Odyssey® Fc imaging system (Li-Cor) using a 30-second exposure.

### Chromatin immunoprecipitation

Hippocampi were extracted as described above and kept at -80°C until further processing. Antibodies were bound and pre-blocked to magnetic beads using the following procedure: fresh block solution was made (225mg bovine serum albumin, 45ml ice cold 1x PBS). Dynabead protein A (50ul; Thermo Fisher 10001D) and protein G (50ul; Thermo Fisher 10003D) were added to 1ml of block solution. Magnetic beads were collected on a magnetic stand for 5min. Block solution was aspirated and this wash cycle was repeated two times. After the final wash, 5ug of anti-CLOCK antibody (Abcam ab3517) or IgG (Abcam ab171870) were added to block solution and a final volume of 250ul was added to the magnetic beads. These beads and antibody solutions were incubated with rotation at 4°C for the duration of chromatin preparation, typically ∼7 hours.

Chromatin preparation and shearing was performed using a Covaris truCHIP chromatin shearing tissue kit (Covaris 520237). Quenching buffer and fixing buffer were made following manufacturer’s instructions (Covaris 520237). Hippocampi (25-40mg) were removed from -80°C and kept on dry ice. Each hippocampus was individually cut to 1mm^3^ segments in a petri dish, kept frozen using liquid nitrogen, then placed in 1.5ml microcentrifuge tubes on dry ice. Tubes containing minced hippocampal tissues were placed at room temperature and 400ul of ice cold 1x PBS was added prior to centrifugation at 4°C for 5-minutes at 3300rpm. During this centrifugation, 1ml of 16% formaldehyde was added to the fixing buffer, to create a final 11.1% formaldehyde fixing buffer. Following centrifugation, supernatant was aspirated and the tissue was placed on ice, resuspended in 400ul of fixing buffer (11.1% formaldehyde), and rocked at room temperature for 8-minutes. Tissue fixation was quenched using 24ul of quenching buffer and incubated at room temperature for 5-minutes while rocking. Tissue was centrifuged at 4°C and 3300rpm for 5-minutes. Supernatant was removed and tissue was washed with 400ul of ice cold 1x PBS and centrifugation at 4°C at 3300rpm for 5min. This 1x PBS wash was repeated twice. After the final wash, supernatant was removed and the tissues were placed in dry ice to flash freeze the tissue.

Tissue pulverization was performed using a Covaris CP02 cryoPREP automated dry pulverizer (Covaris 500001). Freeze dried tissue was placed in Covaris tissue bags (Covaris 520001) and pulverized twice using a setting of 5 on the cryoPREP. Following pulverization, tissue was placed in liquid nitrogen and then moved to dry ice. Lysis buffer, protease inhibitor cocktail, wash buffer, and shearing buffer were made following manufacturer’s instructions (Covaris 520237). Pulverized tissue was thawed on ice and 400ul of lysis buffer was added to each tissue bag to resuspended the tissue with pipetting, and contents were transferred to a new tube on ice. Tubes were rotated at 4°C for 20-minutes, then centrifuged at 4°C for 5-minutes at 2000xg. The supernatant was aspirated and 400ul of wash buffer was added to the tubes before samples were incubated at 4°C for 10-minutes with rotation. Samples were then centrifuged at 4°C for 5-minutes at 2000xg. This wash was repeated two times. Following the final wash, 125ul of shearing buffer was added to tubes and incubated on ice for 10-minutes. Following this incubation, 130ul of the prepared nuclei were transferred to microTUBES (Covaris 520216) and placed on ice. Shearing of chromatin was performed using a Covaris S220 focused-ultrasonicator (500217), using the following parameters: PIP 105, 2% duty factor, CPB 200, treatment time of 8-minutes, setpoint temperature of 6°C, minimum temperature of 3°C, maximum temperature of 9°C, and continuous degassing. Samples were placed on ice following sonication. ChIP dilution buffer was made following manufacturer’s instructions (Covaris 520237). 160ul of ChIP dilution buffer was added to new a 1.5 microcentrifuge tube, followed by the 130ul of sheared chromatin sample. Samples were centrifuged for 10-minutes at 4°C at max speed (20,000xg). Supernatant was transferred to new a 1.5 microcentrifuge tube on ice and protein quantification was performed using a Qubit protein assay kit (Thermo Fisher Q33212) and a Qubit 4 fluorometer (Q33238).

Pre-blocked magnetic bead-bound antibodies were washed in fresh block solution as described above, for an additional three washes. Following the washes, magnetic beads were resuspended in 100ul of block solution. 600ug of protein from sheared samples were placed in corresponding tubes containing anti-CLOCK antibody or IgG control. ChIP dilution buffer was added to a final total volume of 300ul. Immunoprecipitation and IgG samples were incubated overnight at 4°C with rotation. Following overnight incubation, magnetic beads were collected using a magnetic stand for 5-minutes and the supernatant was aspirated. Bead containing tubes were removed from the magnetic stand and resuspended in 1ml of RIPA buffer (50 mM Hepes– KOH, pH 7.5; 500 mM LiCl; 1 mM EDTA; 1% NP-40 or Igepal CA-630; 0.7% Na–Deoxycholate). Magnetic beads were collected on a magnetic stand for 5-minutes and RIPA buffer was aspirated off. This wash cycle was repeated four times. Magnetic beads were resuspended in 1ml of TBS (20 mM Tris–HCl, pH 7.6; 150 mM NaCl) and then collected on a magnetic stand for 5-minutes. TBS was aspirated and tubes were centrifuged at 1000rpm for 3-minutes at 4°C. Tubes were placed on a magnetic stand and any residual TBS was aspirated off. Magnetic beads were resuspended in 200ul of elution buffer (20 mM Tris–HCl, pH 7.6; 150 mM NaCl) and samples were placed in a water bath set at 65°C overnight to reverse-crosslink proteins from DNA.

Following reverse-crosslinking, samples were placed on a magnetic stand for 5min. Supernatant from immunoprecipitated and IgG samples were transferred to new 1.5ml microcentrifuge tubes, and 200ul of TE (10 mM Tris-HCl; pH 8.0; 1mM EDTA) was added, along with 10ug of RNaseA (EN0531), before the sample was incubated in a 37°C water bath for 30-minutes. After the incubation, 4ul of proteinase K (Invitrogen, 25530-049) was added to each sample, mixed, and incubated at 55°C for one hour. 400ul of phenol-chloroform-isoamyl alcohol was added to each tube and mixed before each sample was transferred to a 2ml phase lock gel light tube (FPR5101) and centrifuged for 5-minutes at 16,000xg. The aqueous layer was removed and transferred to a new 1.5ml microcentrifuge tube containing 16ul of 5M NaCl and 1ul of GlycoBlue (AM9516). 800ul of 100% ethanol was added and the contents were mixed and then incubated for 30-minutes at -80°C to precipitate the DNA. Samples were then centrifuged at 4°C for 10-minutes at 20,000xg to pellet the DNA. Pellets were washed with 500ul of 80% ethanol and centrifuged for 5-minutes at 4°C at 20,000xg. Supernatant was aspirated and DNA pellets were allowed to air dry for 30-minutes at room temperature. DNA pellets were resuspended in 50ul of ultra-pure water 10 mM Tris–HCl, pH 8.0 and placed in a 55°C water bath for 10-minutes prior to vortexing.

#### ChIP-sequencing

Next-generation sequence libraries were generated from immunoprecipitated and IgG eluted DNA using the NEBNext ChIP-seq library prep reagent set for Illumina sequencing, according to the manufacturer’s instructions. Briefly, eluted DNA fragments were purified after the adapter ligation step using using AMPure XP beads (Agencourt A63880). An Agilent 2100 Bioanalyzer was used to quantify the amplified library DNA and 20pM of diluted libraries were used for sequencing. A Sage Science Pippin HT was used to size select DNA from libraries, for an average size of ∼400bp per sample. 50-cycle paired-end sequencing was performed by the University of Wisconsin – Madison Biotechnology Center. Image processing and sequence extraction were done using the standard Illumina Pipeline.

#### ChIP-sequencing analysis

Raw paired-end sequencing files were assessed for quality using FASTQC (https://www.bioinformatics.babraham.ac.uk/). Adapters were removed and sequence reads were trimmed, using a quality score cutoff of 30, in the trim_galore package in R (https://www.bioinformatics.babraham.ac.uk/). Trimmed reads were aligned to the *mm9* genome using bowtie2 v2.3.4^91^, local alignment. Following alignment, only uniquely mapped sequence reads were used for downstream analysis, samtools to filter multi-mapping reads^104^ and PCR duplicates. Uniquely-mapped and filtered sequence reads were mapped to differentially hydroxymethylated regions (DhMRs) found to contain a putative CLOCK binding site (CACGTG) by motif enrichment analysis (*N* = 577). Only data from these sites were used for differential binding analyses (R package *edgeR*^93^), after controlling for background through normalization to sequenced IgG reads. A *P*-value threshold of 0.05 was used to determine significant differential binding of CLOCK between groups.

### Electrophoretic mobility shift assays

Complementary 5’-biotinylated (0.5uM) and unlabeled oligonucleotides (Integrated DNA Technologies) were annealed by heating equal concentrations of sense and anti-sense oligonucleotides to 95°C and lowering the temperature by 2°C in 3-minute intervals until reaching 23°C. Mouse hippocampal nuclear extracts were isolated from adult female mice (P90) using a nuclear and cytoplasmic extraction kit per the manufacturer’s instructions (Thermo Scientific 78833). Shift assays were performed using the LightShift® Chemilumiescent EMSA Kit (Thermo Scientific 20148). DNA binding reactions were performed in a 20ul system containing biotinylated oligonucleotides and nuclear extracts (12ug). For cold competition assays, a 1:1000 concentration of unlabeled oligonucleotides was incubated on ice for 1-hour with nuclear extracts prior to the addition on biotinylated oligonucleotides. For supershift assays, 5ug of anti-CLOCK antibody (Abcam, ab3517) or anti-BMAL1 antibody (Abcam, ab93806) were incubated on ice with nuclear extract for 1-hour prior to the addition of biotinylated oligonucleotides. In such assays, nuclear extracts without the addition of unlabeled oligonucleotides or antibody were also incubated on ice for 1-hour prior to the addition of biotinylated oligonucleotides, to ensure the absence of protein degradation during the incubation period. Following incubation on ice, biotinylated oligonucleotides (0.5uM) were added to binding reactions and incubated at room temperature for 20-minutes. Reaction products were separated by electrophoresis at 100V for 60-minutes. Following separation, protein::DNA complexes were transferred onto a positively-charged nylon membrane (Thermo Scientific 77016) for 60-minutes at 380A on ice. Protein::DNA complexes were detected using the Nucleic Acid Detection Module Kit (Thermo Scientific 89880) per the manufacturer’s instructions, and imaged using an Odyssey® Fc imaging system (Li-Cor) and a 60-minute exposure.

### ImageJ band quantification

Band intensity percentages were quantified using ImageJ software. First, images were set to grey-scale. Bands of interest were selected across all lanes and their inverted means were obtained. Background noise intensity was subtracted out by subtracting the intensity of staining above the bands of interest. Band intensities were normalized to loading controls lanes (*i.e.,* beta-actin (western blot)). For EMSA, a two-sided T-test (R environment) was used to determine significant alterations in binding affinity to methylated- and hydroxymethylated-oligonucleotides, using three independent EMSA replicates. For western blot, a two-sided T-test (R environment) was used to determine significant changes in CLOCK expression between groups/genotypes, using three independent immunoblot replicates.

### Oligonucleotide sequences

#### Unmodified Probes

Fry1 FW: 5’-Biotin-TATGTTCATCCACGTGATGC-3’

Fry1 RVL 5’-Biotin-GCATCACGTGGATGAACATA-3’

Gigyf1 FW: 5’-Biotin-GGGTACACGTGCGCCATGGC-3’

Gigyf1 RV: 5’-Biotin-GCCATGGCGCACGTGTACCC-3’

Palld FW: 5’-Biotin-GAATGTCCACACGTGTATGC-3’

Palld RV: 5’-Biotin-GCATACACGTGTGGACATTC-3’

#### 5hmC Probes

Fry1 FW: 5’-Biotin-TATGTTCATCCA^hme^CGTGATGC-3’

Fry1 RV: 5’-Biotin-GCATCA^hme^CGTGGATGAACATA-3’

Gigyf1 FW: 5’-Biotin-GGGTACA^hme^CGTGCGCCATGGC-3’

Gigyf1 RV: 5’-Biotin-GCCATGGCGCA^hme^CGTGTACCC-3’

Palld FW: 5’-Biotin-GAATGTCCACA^hme^CGTGTATGC-3’

Palld RV: 5’-Biotin-GCATACA^hme^CGTGTGGACATTC-3’

#### 5mC Probes

Fry1 FW: 5’-Biotin-TATGTTCATCCA^me^CGTGATGC-3’

Fry1 RV: 5’-Biotin-GCATCA^me^CGTGGATGAACATA-3’

Gigyf1 FW: 5’-Biotin-GGGTACA^me^CGTGCGCCATGGC-3’

Gigyf1 RV: 5’-Biotin-GCCATGGCGCA^me^CGTGTACCC-3’

Palld FW: 5’-Biotin-GAATGTCCACA^me^CGTGTATGC-3’

Palld RV: 5’-Biotin-GCATACA^me^CGTGTGGACATTC-3’

#### Unlabeled Probes

Fry1 FW: 5’-TATGTTCATCCACGTGATGC-3’

Fry1 RVL 5’-GCATCACGTGGATGAACATA-3’

Gigyf1 FW: 5’-GGGTACACGTGCGCCATGGC-3’

Gigyf1 RV: 5’-GCCATGGCGCACGTGTACCC-3’

Palld FW: 5’-GAATGTCCACACGTGTATGC-3’

Palld RV: 5’-GCATACACGTGTGGACATTC-3

## Acknowledgements

The authors would like to thank Dr. Sergio Tufik for his endless support, the UW biotechnology center, and Dr. Sisi Li for her assistance with brain dissections. This work was supported in part by the University of Wisconsin-Madison departments of Neurosurgery and Psychiatry, University of Wisconsin Vilas Cycle Professorships #133AAA2989 and University of Wisconsin Graduate School #MSN184352 (all to RSA), NARSAD Young Investigator Grant from the Brain & Behavioral Research Foundation #22669 & 25212 (LP), Ruth L. Kirschstein National Research Service Award MH113351-02 (AM), University of Wisconsin Neuroscience training grant T32-GM007507 (SL) NIH grants HG003747 and U54AI117924 (all to SK), HG007019 (SK and QZ) and the University of Nebraska Lincoln department of Statistics (QZ). Part of QZ’s research was performed when he was a postdoc in the statistical genomics group at UW Madison led by SK. The authors declare no competing financial interests.

## Author contributions

Conceptualization: LAP, AM, RSA; Methodology: LAP, AM, QZ, KC, LS, SK, RSA; Validation: LAP, AM, LS, RSA; Software: AM, QZ, KC, SK, RSA; Formal analysis: LAP, AM, QZ, KC, RSA; Writing - original draft: LAP, AM, RSA; Writing - review/editing: LAP, AM, QZ, KC, LS, SK, RSA.

## Supplemental Figure Legends

**Supplemental Figure 1:**
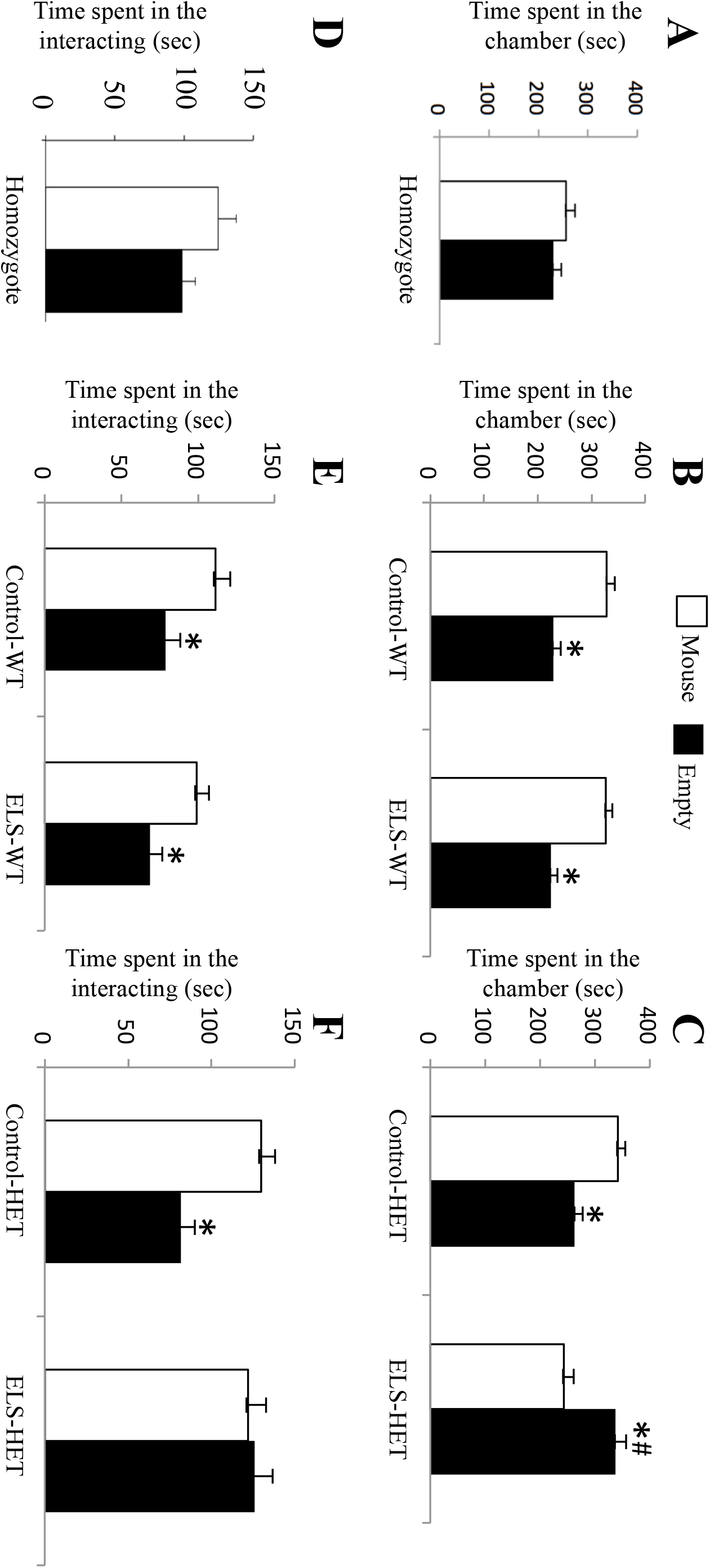
Results from 3-chamber social interaction test in females from an independent cohort. (A-C) test time spent in the chamber with either an unfamiliar mouse inside a cup or with an empty cup. (A) *Cntnap2* homozygote: Data analysis details: Paired T-test did not detect any significant differences. (B) two way ANOVA for <u>control-WT versus ELS-WT</u> <u>groups</u>: effect of time spent in chamber: *F*_(1,16)_ = 58, *P*-value < 0.001. (C) two way ANOVA <u>for control-HET versus ELS-HET groups: interaction between treatment and time spent in chamber</u>: *F*_(1,24)_ = 25, *P*-value < 0.001. There was a significant three-way interaction between effect of treatment, time spent in chamber, and genotype *F*_(1,40)_ = 12.5, *P*-value = 0.001. (D-F) Time spent interacting with either an unfamiliar mouse inside a cup or with an inanimate object. (D) *Cntnap2* homozygote: Data analysis details: Paired T-test did not detect any significant differences. (E) two way ANOVA for <u>control-WT versus ELS-WT groups</u>: effect of time spent interacting: *F*_(1,16)_ = 12.3, *P*-value = 0.003. (F) two way ANOVA <u>for control-HET versus ELS-HET groups:</u> effect of time spent interacting *F*_(1,24)_ = 4.7, *P*-value = 0.004 and interaction between treatment and time spent interacting *F*_(1,24)_ = 6.5, *P*-value = 0.01. The asterisk (*) denotes significance within group (*P*-value < 0.05). Hashtag (#) denotes significance for time spent in empty chamber between ELS-HET in comparison to Control-WT, Control-HET, and ELS-WT (*P*-value < 0.05).

**Supplemental Figure 2:**
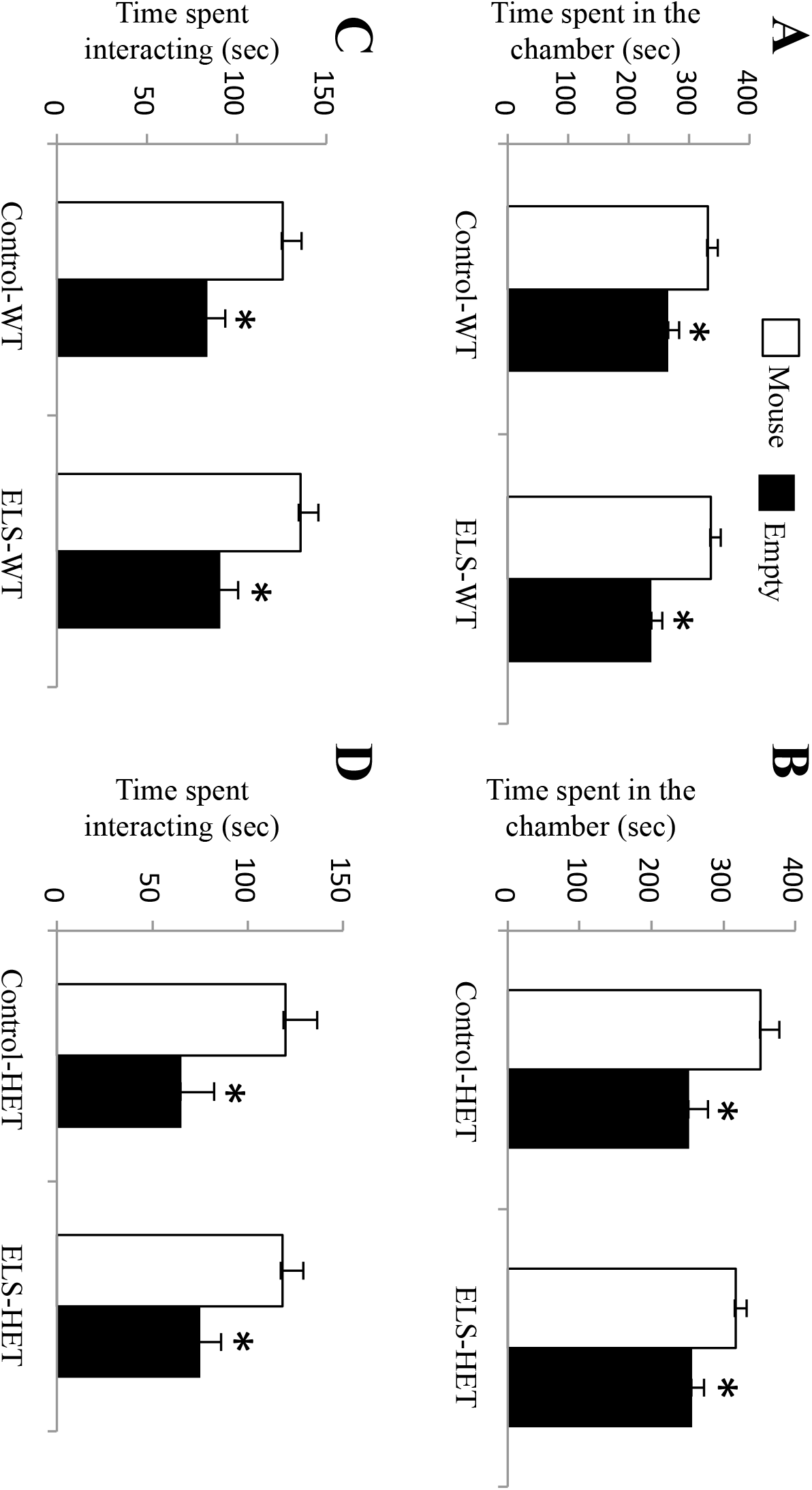
Results from social behavioral tests in male offspring. (A-B) 3-chamber social interaction test, time spent in the chamber with either an unfamiliar mouse inside a cup or with an empty cup. (A) two way ANOVA for <u>control-WT versus ELS-WT groups</u>: effect of time spent in chamber: *F*_(1,28)_ = 22.4, *P*-value < 0.001. (B) two way ANOVA <u>for</u> <u>control-HET versus ELS-HET groups:</u> effect of time spent in chamber: *F*_(1,24)_ = 13.1, *P*-value = 0.001. (C-D) Time spent interacting with either an unfamiliar mouse inside a cup or with an empty cup. (C) two way ANOVA for <u>control-WT versus ELS-WT groups</u>: effect of time spent in chamber: *F*_(1,28)_ = 18.5, *P*-value < 0.001. (D) two way ANOVA <u>for control-HET versus ELS-HET groups:</u> effect of time spent in chamber: *F*_(1,24)_ = 10.7, *P*-value = 0.003. The asterisk (*) denotes significance of *P*-value < 0.05.

**Supplemental Figure 3:**
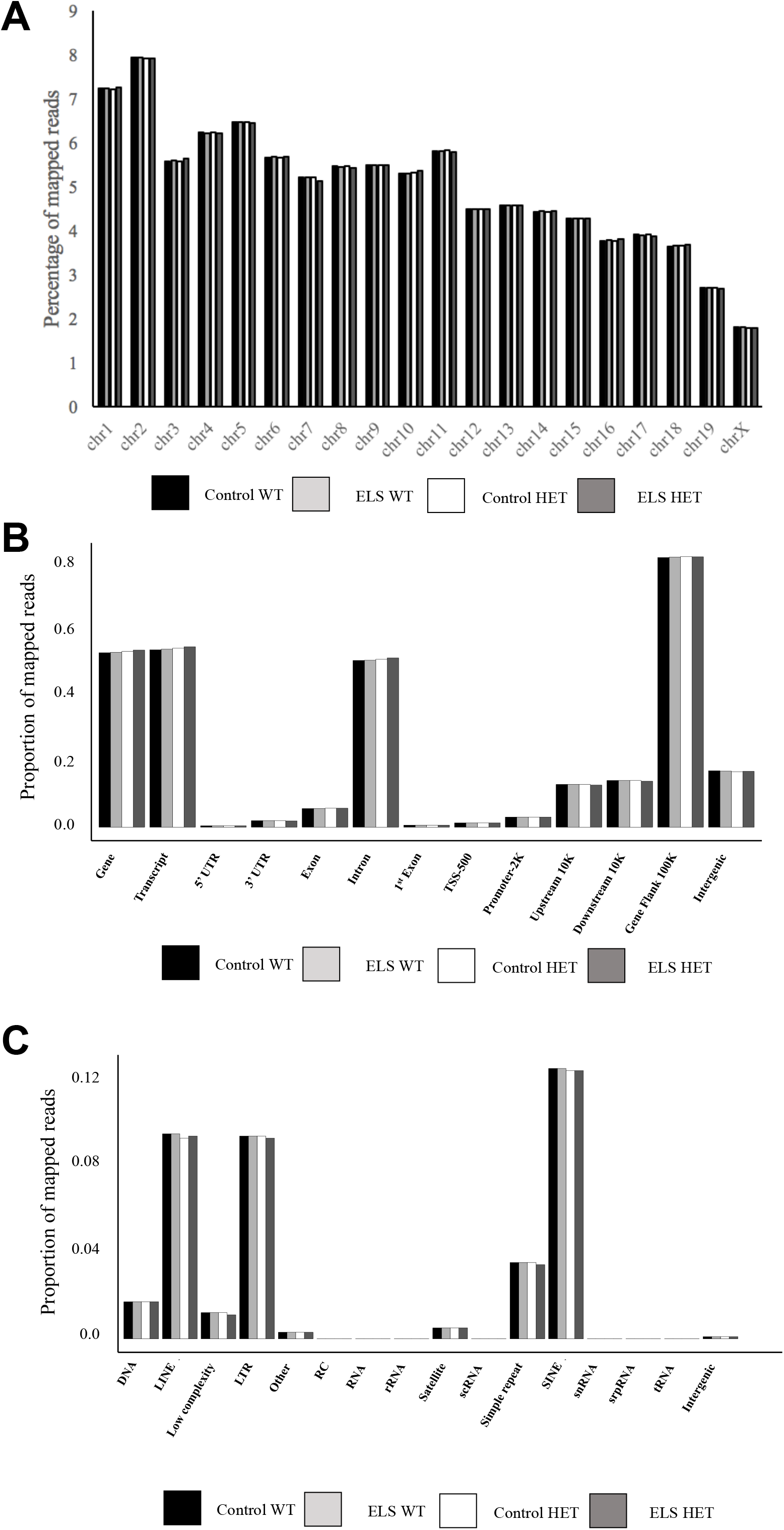
Distribution and characterization of aligned reads derived from hippocampal tissue from the four experimental groups. (A) A bar plot depicts the percent of mapped reads (y-axis) relative to chromosomes (x-axis) for Control-WT (black bars), ELS-WT (light grey bars), Control-HET (white bars), and ELS-HET (dark grey bars) mice. (B) A bar plot shows the proportion of mapped reads (y-axis) as they relate to standard genomic structures (x-axis) for Control-WT (black bars), ELS-WT (light grey bars), Control-HET (white bars), and ELS-HET (dark grey bars) mice. (C) A bar plot shows the proportion of mapped reads (y-axis) as they relate to repetitive elements across the genome (x-axis) for Control-WT (black bars), ELS-WT (light grey bars), Control-HET (white bars), and ELS-HET (dark grey bars) mice.

**Supplemental Figure 4:**
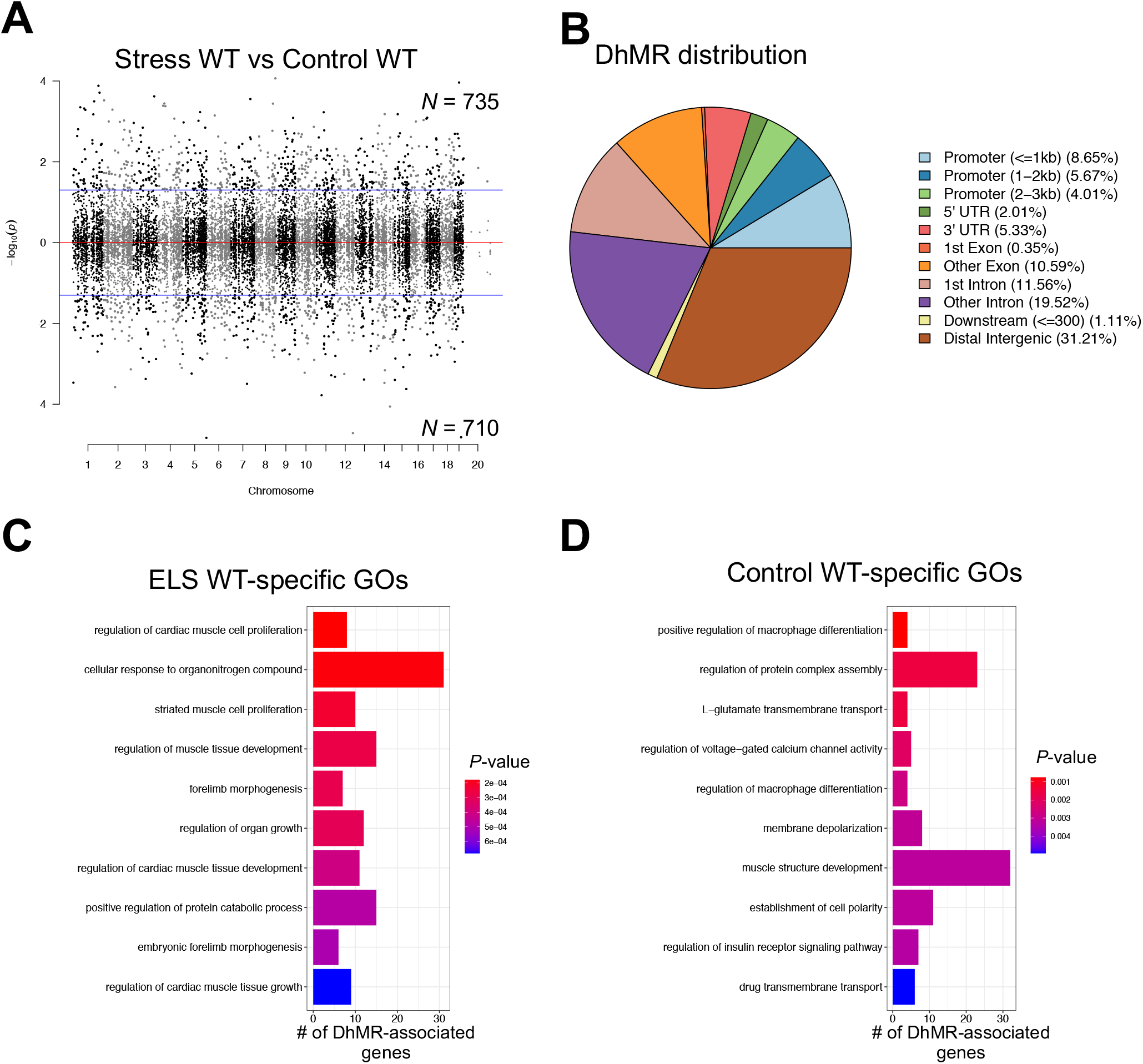
Distribution of hippocampal WT DhMRs. (A) A Manhattan plot shows the distribution of 5hmC sequenced data across the genome (x-axis; alternating black/grey for alternating chromosomes) and the level of significance (y-axis; -log10(*P*-value)). Dots above the top blue line represent ELS-WT-specific-(hyper) DhMRs, while dots below the bottom blue line represent Control-WT-specific-(hypo) DhMRs (*P*-value < 0.05). (B) A pie chart displays the proportion of WT-DhMRs across standard genomic structures. (C-D) Bar plots showing the top ten significantly over-represented ontological terms (y-axis) based on gene ontological analysis of ELS-WT-specific DhMR-associated genes (C) and Control-WT-specific DhMR-associated genes (D). The number of DhMR-associated genes linked to the ontological terms are displayed on the x-axis. Bar color is based on *P*-value.

**Supplemental Figure 5:**
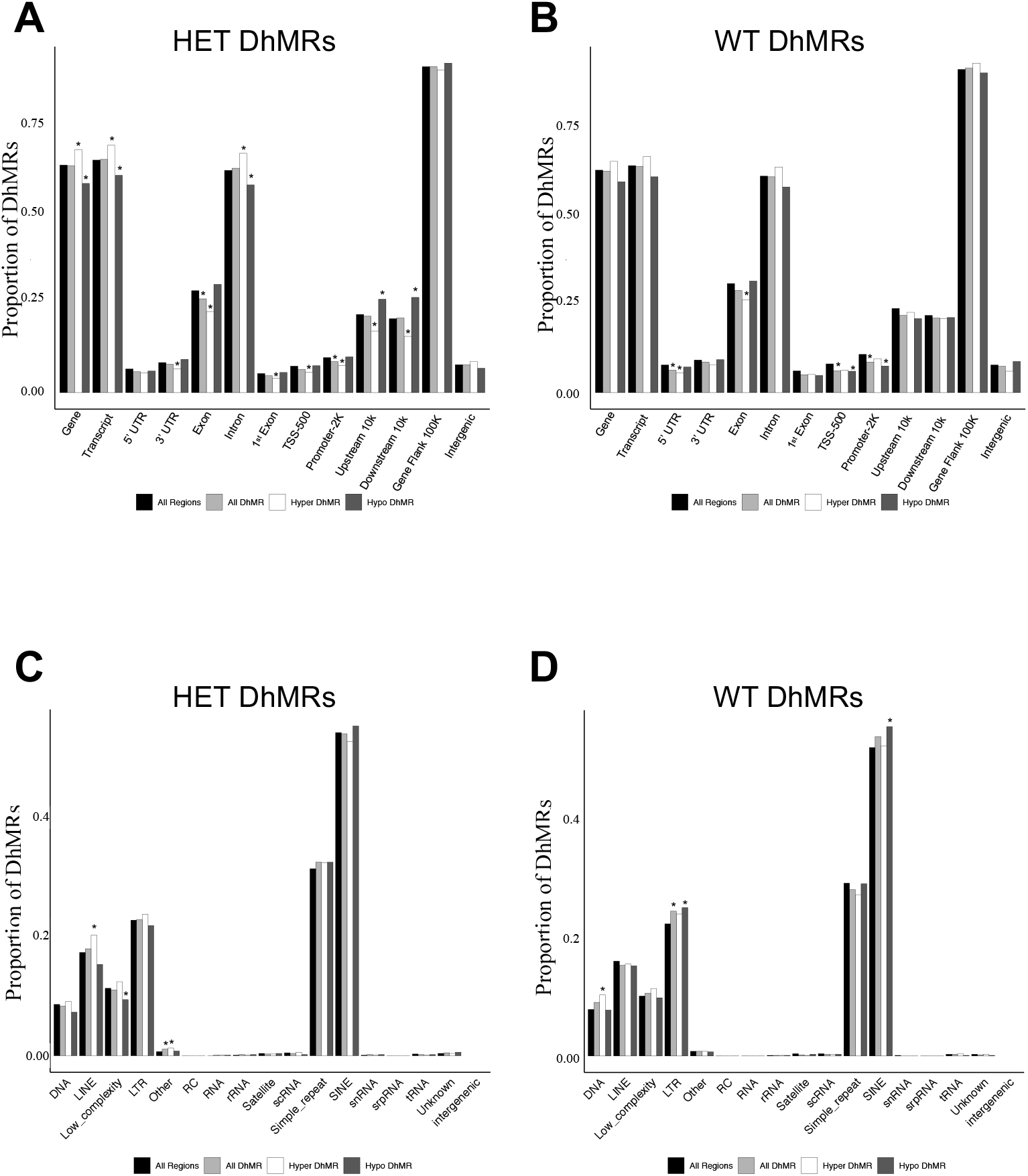
Distribution of hippocampal DhMRs. (A and B) Bar plots showing the proportion of DhMRs (y-axis) from the ELS-HET vs Control-HET (A) and ELS-WT vs Control-WT (B) comparisons as they relate to standard genomic structures (x-axis). Depicted are all regions across the genome tested for differential 5hmC abundance (*i.e.,* background regions; black bars), all DhMRs (light grey bars), ELS-specific (hyper) DhMRs (white bars), and Control-specific (hypo) DhMRs (dark grey bars). An asterisk (*) represents a significant increase or decrease of 5hmC abundance relative to the background regions, as determined by binomial testing (*P*-value < 0.05). (C and D) Bar plots showing the proportion of DhMRs (y-axis) from the ELS-HET vs Control-HET (C) and ELS-WT vs Control-WT (D) comparisons as they relate to repetitive elements (x-axis). Depicted are all regions across the genome tested for differential 5hmC abundance (*i.e.,* background regions; black bars), all DhMRs (light grey bars), ELS-specific (hyper) DhMRs (white bars), and Control-specific (hypo) DhMRs (dark grey bars). An asterisk (*) represents a significant increase or decrease of 5hmC disruptions relative to the background regions, as determined by binomial testing (*P*-value < 0.05).

**Supplemental Figure 6:**
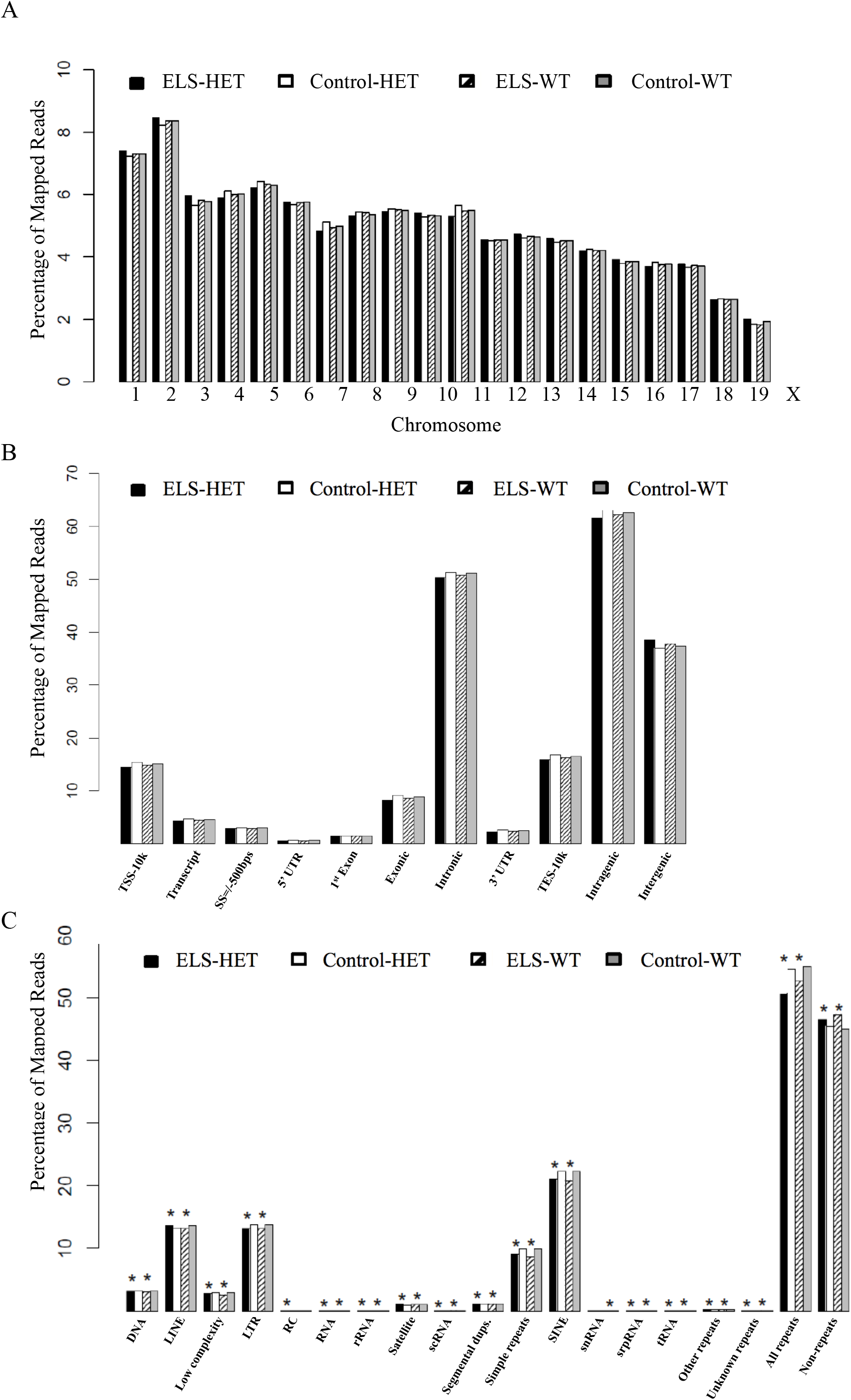
Distribution and characterization of aligned reads derived from striatal tissue from the four experimental groups. (A) A bar plot depicts the percent of mapped reads (y-axis) relative to chromosomes (x-axis) for ELS-HET (black bars), Control-HET (white bars), ELS-WT (striped bars), and Control-WT (grey bars) mice. (B) A bar plot shows the percent of mapped reads (y-axis) as they relate to standard genomic structures (x-axis) for ELS-HET (black bars), Control-HET (white bars), ELS-WT (striped bars), and Control-WT (grey bars) mice. (C) A bar plot shows the proportion of mapped reads (y-axis) as they relate to repetitive elements across the genome (x-axis) for for ELS-HET (black bars), Control-HET (white bars), ELS-WT (striped bars), and Control-WT (dark grey bars) mice. An asterisk (*) represents a significant increase or decrease of 5hmC disruptions relative to the background regions, as determined by binomial testing (*P*-value < 0.05).

**Supplemental Figure 7:**
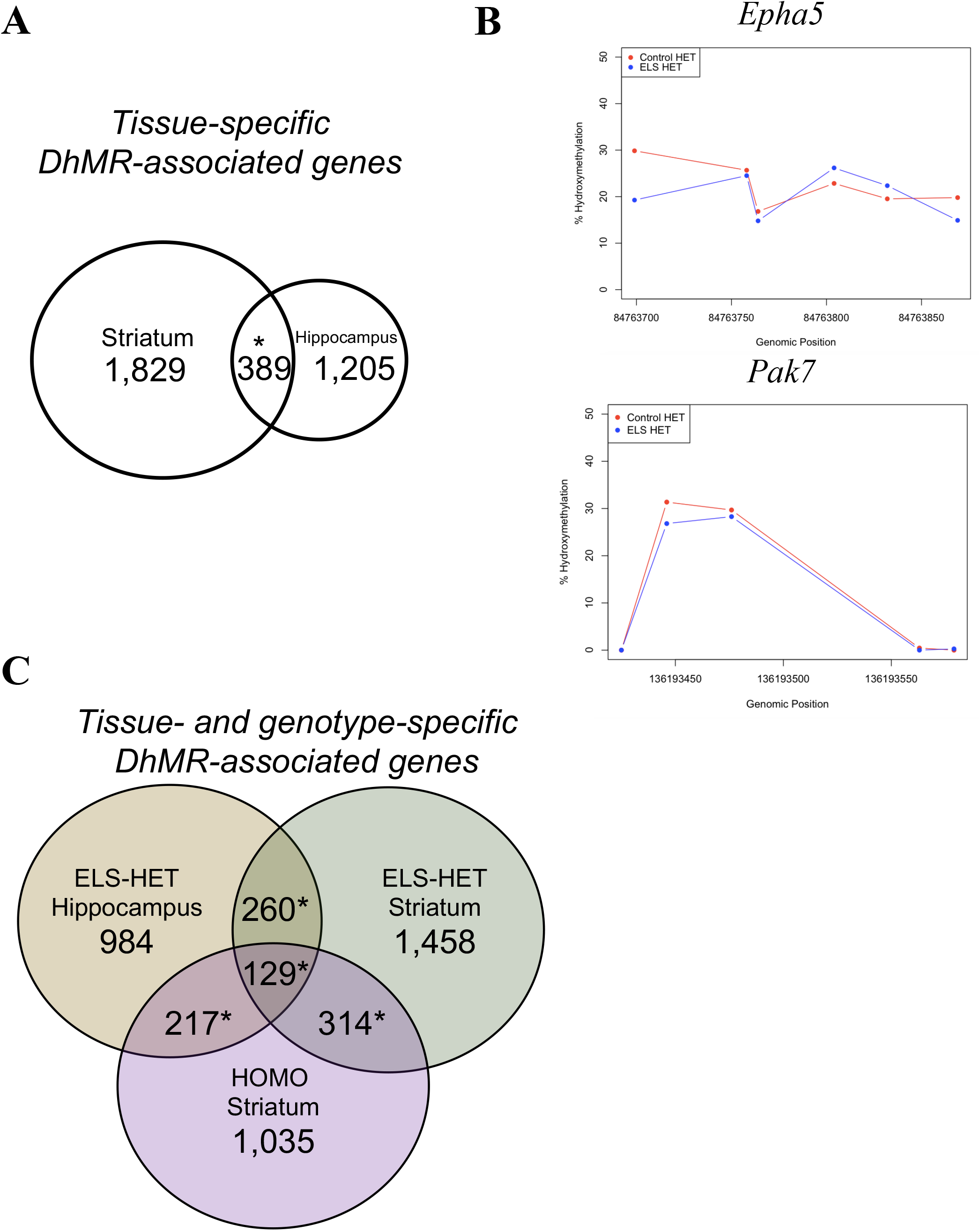
Molecular validation of hippocampal DhMR data. (A) A Venn diagram displays the overlap of DhMR-associated genes found in hippocampal and striatal tissues. An asterisk (*) represents a significant enrichment of DhMR-associated genes between the two tissues, as determined by a hypergeometric enrichment test (*P*-value < 0.05). (B) Molecular validation of a DhMR (*Epha5*) and a non-DhMR (*Pak7*). The genomic position (x-axis) and percent hydroxymethylation (y-axis) as determined by simultaneous targeted methylation sequencing of hippocampal tissue from ELS-HET (blue) and Control-HET (red) mice. An asterisk (*) represents a significant increase or decrease of 5hmC abundance of ELS-HET mice compared to Control-HET mice, as determined by Fisher’s exact testing (*P*-value < 0.05). (C) A Venn diagram shows the overlap of DhMR-associated genes found in hippocampal and striatal tissues from ELS *Cntnap2^+/-^* (ELS-HET) and *Cntnap2^-/-^* (HOMO) mice. An asterisk (*) represents a significant enrichment of DhMR-associated genes between the two tissues, as determined by a hypergeometric enrichment test (*P*-value < 0.05).

**Supplemental Figure 8:**
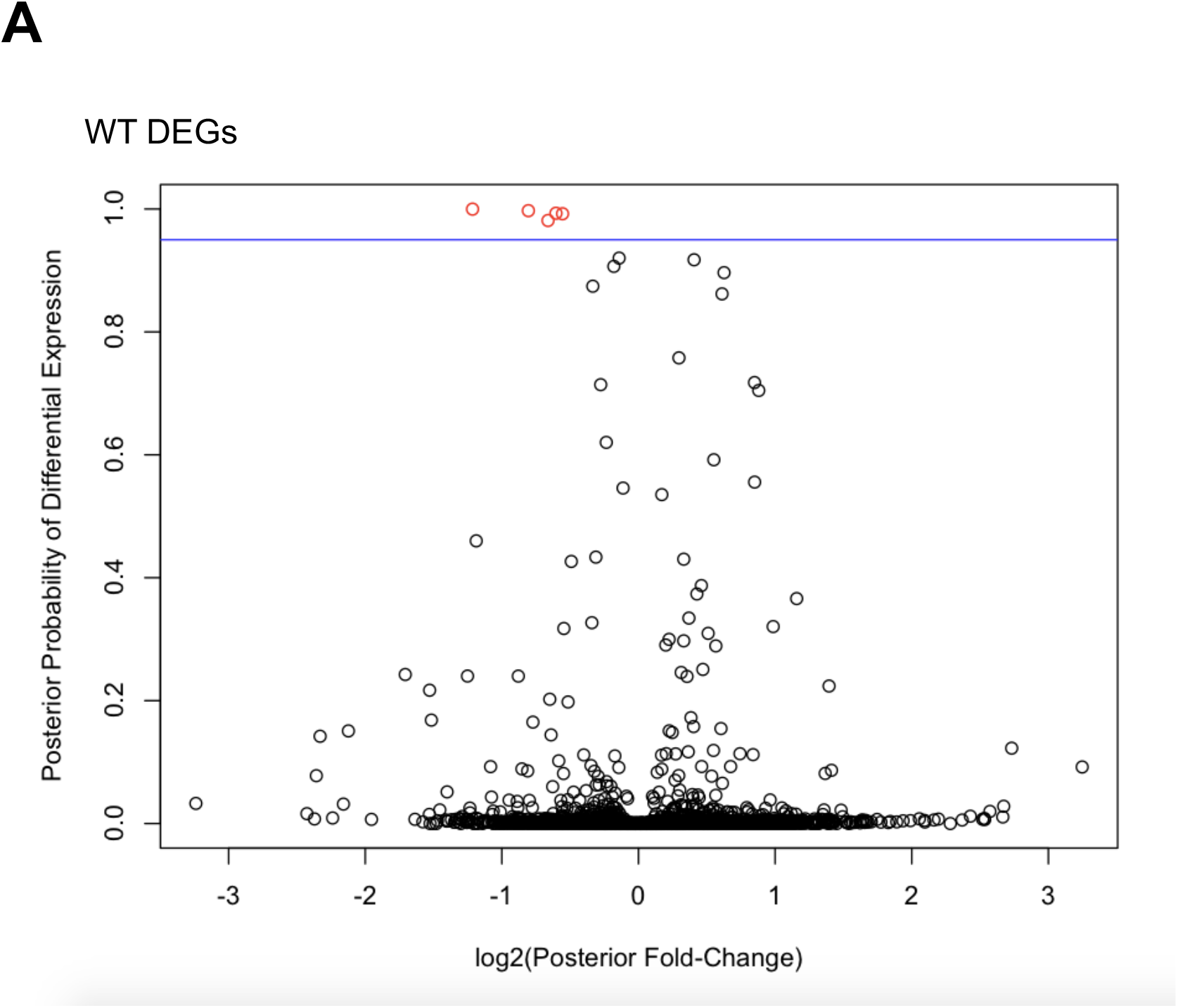
Characterization of differentially expressed genes (DEGs) between ELS-WT and Control-WT. (A) A modified volcano plot depicts the log2(posterior fold change; x-axis) versus the posterior probability of differential expression (y-axis). Open circles (red/black) represent each gene examined and differentially expressed genes (red) are shown above the significance line (blue, FDR *P*-value < 0.1).

**Supplemental Figure 9:**
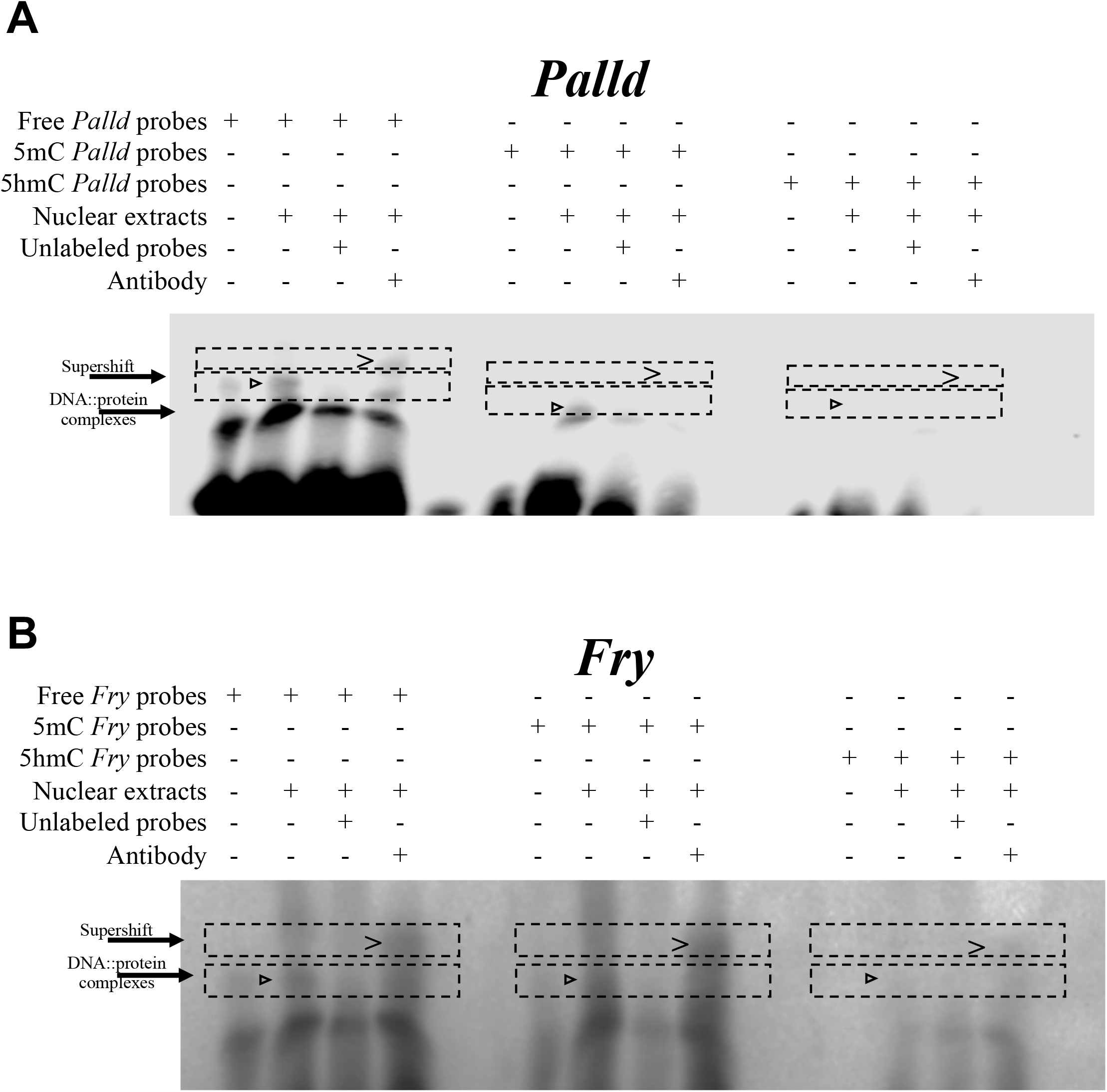
Functional assays reveals that 5hmC represses CLOCK binding. (A-B) Electromobility shift assays with hippocampal lysates. Lysate shift (lanes 2, 6, and 10), cold competition (lanes 3, 7, and 11), and supershift (lanes 4, 8, and 12) of DNA probes corresponding to a CLOCK E-box motif in the DhMR of *Palld* (A) and *Fry* (B) that exhibited differential CLOCK binding. Biotinylated DNA probes contained either an unmodified E-box motif (lanes 1-4), a methylated E-box motif (lanes 5-8), or a hydroxymethylated E-box motif (lanes 9-12). Regions containing bands of interest are highlighted (dashed boxes) with DNA::protein complexes that are abrogated with cold competition are indicated (black arrowheads) and supershift complexes are shown (carrot).

**Supplemental Table 1:** Summary of behavioral tests in male offspring

|  | **Control** |  | **Early-Life Stress** |  |
| --- | --- | --- | --- | --- |
|  | **WT** | ***Cntnap2*<sup>+/-</sup>** | **WT** | ***Cntnap2*<sup>+/-</sup>** |
| **Open Field** |  |  |  |  |
| Time spent in the center (s) | 47 ± 6 | 35 ± 7 | 42 ± 4 | 54 ± 6 |
| Number of entries to center | 19 ± 3 | 14 ± 1 | 19 ± 2 | 20.9 ± 2 |
| **Elevated Plus Maze** |  |  |  |  |
| Time spent in closed arm (s) | 209 ± 13 | 213 ± 13 | 213 ± 12 | 214 ± 8 |
| Time spent in open arm (s) | 204 ± 16 | 208 ± 10 | 202 ± 10 | 294 ± 8 |
| **Light/Dark Box** |  |  |  |  |
| Time spent in light side (s) | 386 ± 14 | 378 ± 25 | 381 ± 12 | 376 ± 17 |
| **Forced swim test** |  |  |  |  |
| Time spent floating | 157 ± 8 | 156 ± 11 | 145 ± 9 | 155 ± 8 |

Comparison of different behavioral tests between early-life stress (ELS) and non-stressed (Control) *Cntnap2*<sup>+/-</sup> and WT mice. The value are presented as mean ± S.E.M. \* $p \leq 0.05$ . N = 8-10. Data analysis: 1) Open Field (time spent in the center: effect of ELS $F_{(1,32)} = 1.06$ , $P = 0.3$ ; effect of genotype $F_{(1,32)} = 0.0009$ , $P = 0.97$ ; number of entries to the center: effect of ELS $F_{(1,32)} = 1.28$ , $P = 0.26$ ; effect of genotype $F_{(1,32)} = 0.29$ , $P = 0.59$ ), 2) Elevated Plus Maze (time spent in closed arm: effect of ELS $F_{(1,35)} = 0.04$ , $P = 0.82$ ; effect of genotype $F_{(1,35)} = 0.04$ , $P = 0.82$ ; time spent in open arm: effect of ELS $F_{(1,35)} = 0.5$ , $P = 0.48$ ; effect of genotype $F_{(1,35)} = 0.02$ , $P = 0.87$ ), 3) Light/Dark Box (time spent in light side: effect of ELS $F_{(1,30)} = 0.03$ , $P = 0.86$ ; effect of genotype $F_{(1,30)} = 0.14$ , $P = 0.7$ ) and 4) Forced Swim Test (time spent floating: effect of ELS $F_{(1,35)} = 0.42$ , $P = 0.5$ ; effect of genotype $F_{(1,35)} = 0.23$ , $P = 0.63$ ).

**Supplementary Table 2:** Hippocampal DNA sequence read information for each biological replicate pool

| Replicates | Total Reads | Monoclonal Reads |
| --- | --- | --- |
| H1_Control_WT | 31009917 | 28354228 |
| H2_Control_WT | 34141544 | 31281271 |
| H3_Control_WT | 27708730 | 25349461 |
| H4_Stess_HET | 31616944 | 29011105 |
| H5_Stress_HET | 30770924 | 28164679 |
| H6_Stress_HET | 32608364 | 29846521 |
| H7_Stress_WT | 35939256 | 32905840 |
| H8_Stress_WT | 35131002 | 31950304 |
| H9_Stress_WT | 25443415 | 23313623 |
| H10_Control_HET | 47224464 | 43340378 |
| H11_Control_HET | 29163771 | 26757595 |
| H12_Control_HET | 28371476 | 25917067 |

**Supplemental Table 3:**
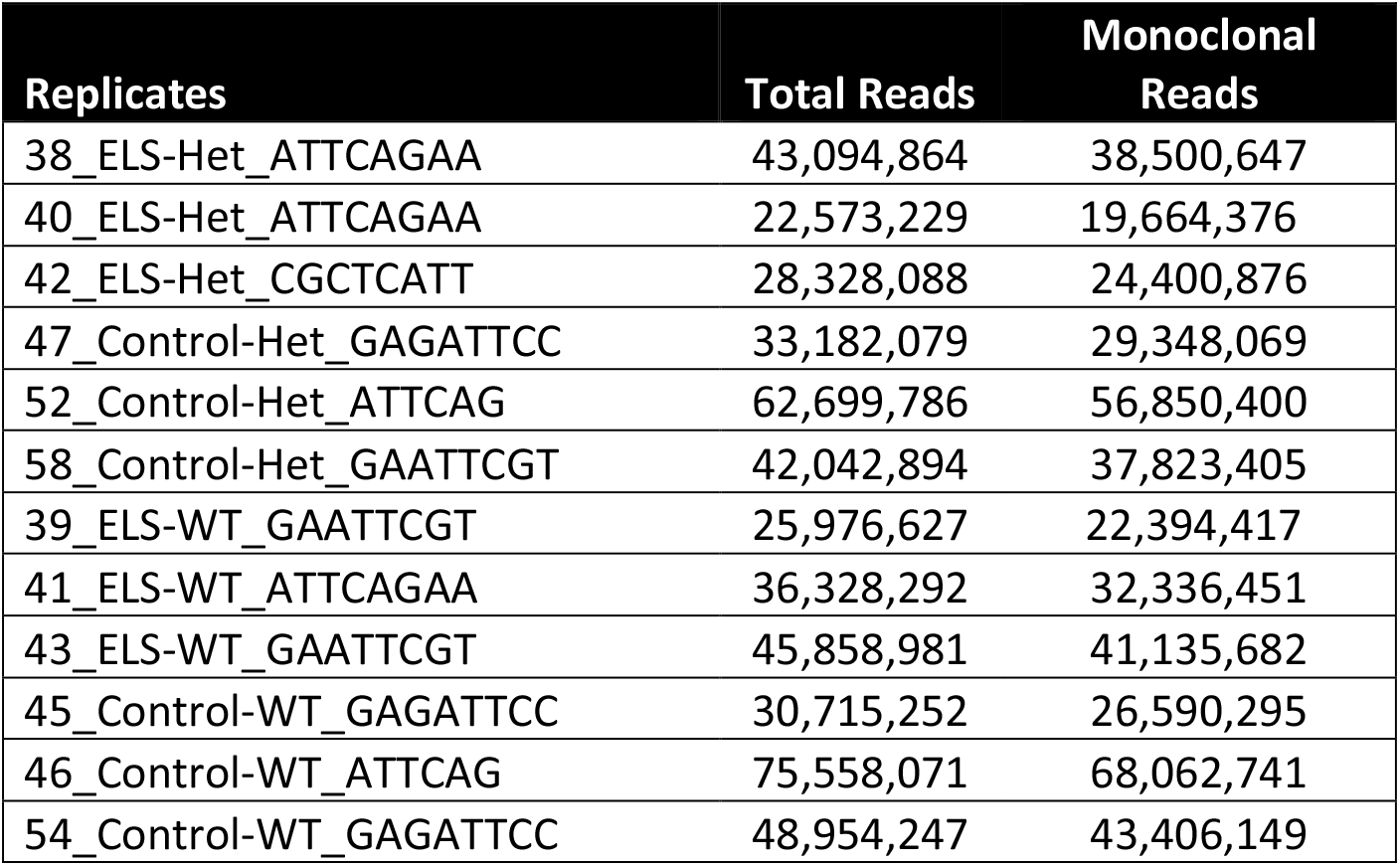
Striatal DNA sequence read information for each biological replicate

| Replicates | Total Reads | Monoclonal Reads |
| --- | --- | --- |
| 38_ELS-Het_ATTCAGAA | 43,094,864 | 38,500,647 |
| 40_ELS-Het_ATTCAGAA | 22,573,229 | 19,664,376 |
| 42_ELS-Het_CGCTCATT | 28,328,088 | 24,400,876 |
| 47_Control-Het_GAGATTCC | 33,182,079 | 29,348,069 |
| 52_Control-Het_ATTCAG | 62,699,786 | 56,850,400 |
| 58_Control-Het_GAATTCGT | 42,042,894 | 37,823,405 |
| 39_ELS-WT_GAATTCGT | 25,976,627 | 22,394,417 |
| 41_ELS-WT_ATTCAGAA | 36,328,292 | 32,336,451 |
| 43_ELS-WT_GAATTCGT | 45,858,981 | 41,135,682 |
| 45_Control-WT_GAGATTCC | 30,715,252 | 26,590,295 |
| 46_Control-WT_ATTCAG | 75,558,071 | 68,062,741 |
| 54_Control-WT_GAGATTCC | 48,954,247 | 43,406,149 |

